# Circadian clock components CCA1 and LHY shape heat responses across regulatory layers

**DOI:** 10.64898/2026.11.19.767892

**Authors:** Rachel I Strout, Lauren E Grubb, Richard Glen Uhrig, Dawn H. Nagel

**Affiliations:** Department of Botany and Plant Sciences, University of California, Riverside, Riverside, California, USA; Department of Biological Sciences, University of Alberta, Edmonton, AB, Canada; Department of Biochemistry, University of Alberta, Edmonton

**Keywords:** Gating, heat-response, circadian clock, non-cycling, CCA1, LHY

## Abstract

**Background:** High temperatures reprogram gene regulatory responses and are particularly deleterious for crop yield, especially when these events occur at night. This emphasizes that partitioning of heat responses is based on time of day. Coordination between time of day and environmental responses relies on the circadian clock. However, the precise regulatory mechanism of how the clock achieves this partitioning at different levels of gene regulation is poorly understood.

**Results:** Using circadian clock mutants with altered *CCA1* and *LHY* expression, we generated a time-resolved transcriptome, proteome, and metabolome following a short-term heat stress in Arabidopsis. We identified regulatory connections specific to CCA1 and LHY, as well as those shared with other clock components linked to heat stress responses. Additionally, new connections between CCA1 and LHY and non-cycling genes highlight a broader role for the clock in regulating heat responsive gene expression. CCA1 and LHY regulated processes, such as sulfur assimilation and ROS scavenging, may contribute to time-of-day specific thermotolerance. Overall, heat responses were dampened in both the transcriptome and proteome in *cca1lhy*, suggesting both transcriptional and post-translational circadian-regulated processes are important for heat responses. Lastly, we identified differences in metabolic responses in *cca1lhy* and suggest a role for circadian regulation of starch degradation in thermotolerance.

**Conclusions:** This study generates and utilizes data across levels of gene regulation to identify heat responses in the transcriptome and proteome regulated by CCA1 and LHY. These findings establish shared and distinct regulatory mechanisms of clock components in response to heat stress and provide new candidates for network integration and a foundation to engineer plants more tolerant to high temperatures depending on time of day.

## Background

Circadian alignment with the environment is important for ensuring key biological processes occur at time of day or year when it is most beneficial [1,2]. To coordinate these daily and seasonal responses, plants integrate cycles of light, temperature, and metabolite levels to entrain the circadian clock [3–5]. The core clock of *Arabidopsis thaliana* (Arabidopsis) consists of the morning expressed transcription factors (TFs) CIRCADIAN CLOCK ASSOCIATED 1 (CCA1) and LATE ENLONGATED HYPOCOTYL (LHY) which repress and are repressed by the evening expressed TIMING OF CAB EXPRESSION 1 (TOC1) [6,7]. In addition, CCA1 and LHY regulate multiple other circadian clock components throughout the 24 h period [8–10].

As morning expressed genes, CCA1 and LHY have been shown to regulate a large proportion of genes that show peak expression in the evening [11–13]. However, genes expressed in the morning and to a lesser extent other times of day are also regulated by CCA1 and LHY, with an estimate based on genome-wide DNA binding and transcriptome data that in total CCA1 and LHY regulate thousands of genes [11–15]. Interaction with target genes is primarily facilitated through the evening element (AAATATCT) and CCA1 binding site (CBS; AAAAATCT) [11,13,16]. However, additional enriched elements are present in genes occupied by CCA1 and LHY suggesting they may be recruited to target genes via other proteins [11–13,17]. Notably, previous studies have also shown that CCA1 and LHY occupy the promoters of not only cycling genes but also non-cycling genes in both diel and circadian conditions highlighting a broader role for their function outside of clock regulation [11–13].

The relationship between the circadian clock and temperature depends on the extremity of the temperature. Temperatures within physiologically relevant ranges (17°C-27°C) can entrain the clock but may also be buffered from altering clock period through temperature compensation mechanisms [18,19]. Temperature increases that may be considered stressful (>32°C), can alter circadian timing and clock gene expression [14,20]. For example, under high temperatures (37°C), the mRNA abundance of *CCA1* is upregulated, while *LHY* is downregulated [14,21].

High temperatures also increase the frequency of a partial retention of the fourth intron of CCA1 (CCA1βb) [22–26]. While less is known about heat induced alternative splicing of LHY, cooler temperatures impact differential isoform accumulation [25,27]. *CCA1* may also be post-transcriptionally regulated where it is less upregulated by heat in the translatome compared to the transcriptome [21]. Additionally, temperature specific phosphorylation of CCA1 by CASEIN KINASE 2 (CK2) appears to antagonize the ability of CCA1 to bind to DNA [28]. While this interaction has been suggested to contribute to temperature compensation of the clock at high ambient temperatures, CK2 and CCA1 interactions have not been examined under heat stress conditions.

How the clock network coordinates the extensive control of gene regulation in response to heat stress likely involves multiple mechanisms to ensure optimal thermotolerance. In Arabidopsis, post-transcriptional processes including alternative splicing, mRNA stability, mRNA sequestration into biomolecular condensates, translation, and post-translational modifications are largely altered in response to heat stress [29–33]. Many of these processes are also regulated by the clock [21,34–38]. For example, the clock component ZEITLUPE (ZTL), an E3 ubiquitin ligase, is involved in the degradation of aggregated proteins in response to high temperature [20].

The clock also regulates the magnitude of responses to temperature stress in a time-of-day specific manner, termed circadian gating [39–44]. This is suggested to ensure that both the anticipation and response to a stress is appropriately timed throughout the day. Specific clock components have been implicated in the gating of temperature responses. For example, CCA1 and LHY, as well as additional clock components such as the PSEUDO RESPONSE REGULATORS (PRRs), REVEILLE 4 (RVE4), RVE8, NIGHT LIGHT INDUCIBLE AND CLOCK-REGULATED GENE 3 (LNK3), and LNK4 have been shown to gate cold responses [41,45,46]. We found in our recent study that approximately one third of the heat-responsive translatome is gated [21]. We also identified that *CCA1* and *LHY* alter temperature responses in the morning, as thousands of genes differ in their response to heat between WT and *cca1lhy* [14]. However, this study was limited to a single time point and performed under diel conditions, limiting a comprehensive understanding of the scope of CCA1 and LHY coordination of heat responsive gene expression throughout the day. Furthermore, studies in crops have revealed that warm night-time temperatures contribute to dramatic reduction in yields and reprogramming of the transcriptome [47]. It has been suggested that the impact of nighttime heat stress may be distinct or controlled by different mechanisms, highlighting the importance of investigating thermo-responsive regulatory mechanisms mediated by the clock throughout the day [42,43,47,48].

It has been proposed that the clock fine-tunes transcriptional heat responses in both a HEAT SHOCK FACTOR (HSF) dependent and independent manner. Multiple HSF and HEAT SHOCK PROTEINS (HSPs) are rhythmic and bound by clock components, suggesting that their regulation of heat responses is likely circadian controlled [49]. Additionally, the clock components RVE4, RVE8, and their protein interactors LNK1 and LNK2 have been demonstrated to regulate early heat responses in an HSF-independent manner [50]. However, whether the core clock components CCA1 and LHY regulate specific or integrated heat stress responsive pathways to fine-tune thermotolerance at different times of day remains poorly understood.

To address this gap in knowledge, we performed time-resolved or time-specific transcriptomics, proteomics and metabolomics following exposure to a short term (1 h 37°C) heat-stress in Arabidopsis seedlings mis-expressing *CCA1* and *LHY* under circadian conditions. We found that transcriptional gating of heat responses is generally lost in *cca1lhy,* and our results suggest that CCA1 and LHY may regulate heat-responsive transcription through both HSF-dependent and -independent mechanisms. Specific and shared targets with other clock components previously linked to heat stress gating implicates both a partitioned and interconnected temporal network underlying thermotolerance in Arabidopsis. We identified several rhythmic and non-rhythmic CCA1 and LHY targets that highlight new regulatory links and pathways that may be important in coordinating heat responses such as ROS scavenging, sulfur assimilation, and starch degradation. Notably, the magnitude of heat responses is generally reduced in *cca1lhy* in the transcriptome, a pattern that persists in the proteome. By comparing between the transcriptome and proteome, we hypothesize a role for circadian regulation of post-transcriptional heat-responses. Additionally, we identify metabolites altered in *cca1lhy* after heat stress and suggest that circadian regulation of starch metabolism may contribute to thermotolerance.

## Results

### CCA1 and LHY reshape the transcriptional landscape of cycling genes

To first determine the extent of CCA1 and LHY mediated perturbation of the transcriptional landscape throughout the day, we performed a time-resolved transcriptomic analysis in wild-type (WT) and *cca1lhy* mutant Arabidopsis seedlings. Briefly, seedlings were grown for 10 days in 12 h light: 12 h dark cycles at constant ambient (22°C) temperatures and then transferred to constant light (LL) for 2 days. On the third day in LL, samples were collected every 3 hours (h) starting at subjective lights ON (ZT48) and RNA-sequencing was performed (Fig. 1a). Differences in genotype was reflected by 13% of the variation between samples (Fig. S1a). Overall, the influence of time on the variation between samples is largely disrupted in *cca1lhy* compared to WT, highlighting the impact of circadian dysregulation (Fig. S1b and S1c). Consistent with previous studies, the mRNA abundance of major components of the circadian oscillator that are regulated by CCA1 and LHY such as *TOC1, ELF3, ELF4,* and *LUX* are induced in *cca1lhy*, while *PRR9* and *PRR7* are repressed (Fig. S2a) [9].

**Figure 1:**
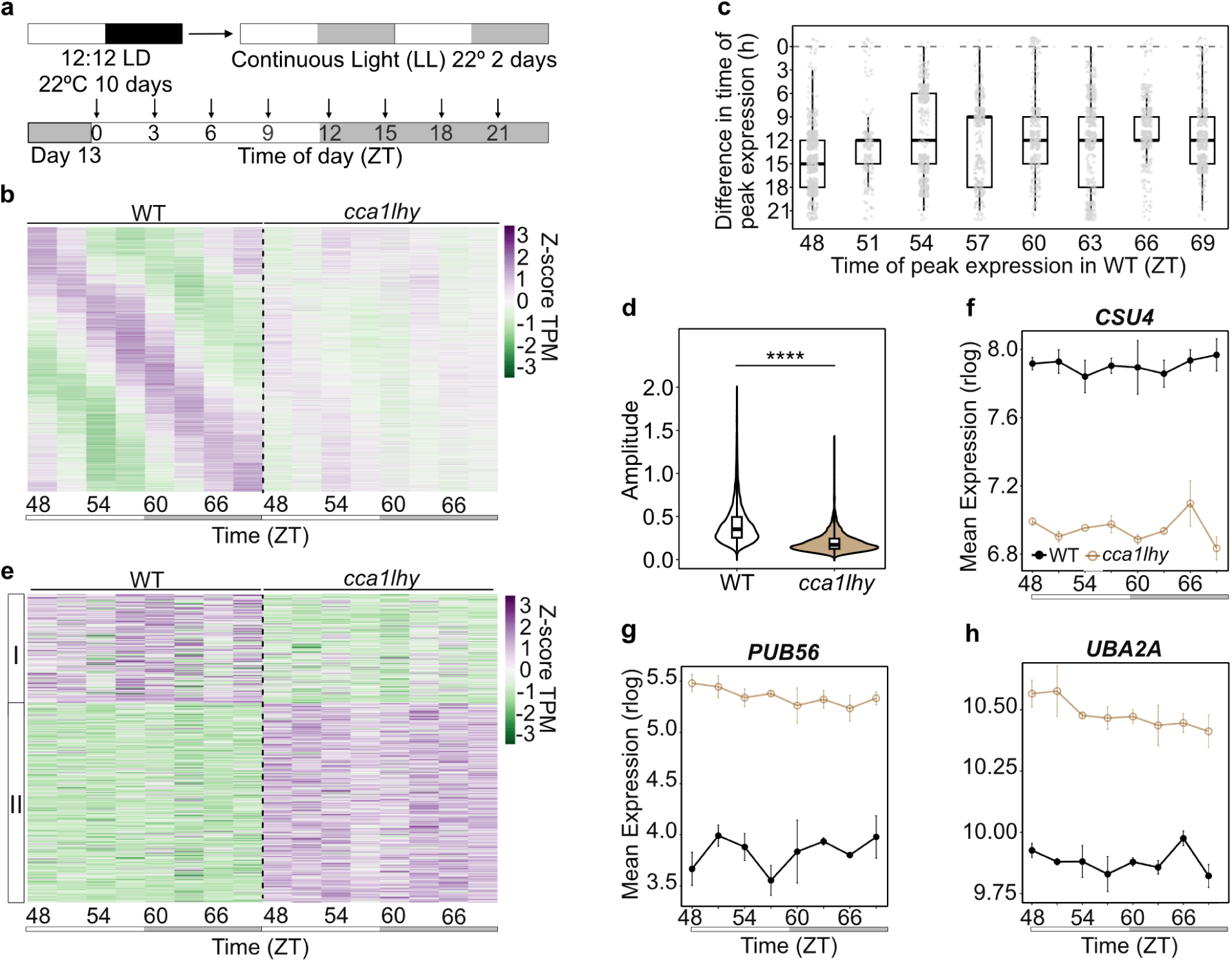
CCA1 and LHY regulate cycling and non-cycling genes. **a)** Experimental design. **b)** The 3101 genes significantly cycling in WT plotted across both genotypes. Columns represent the average of replicates (N=2 or 3). Cycling genes were determined using Metacycle BH. Q < 0.05. Data is scaled by row and ordered by timing of peak abundance. **c)** The difference in time of peak expression in *cca1lhy* of the genes that cycle in WT. Genes separated by the time of peak expression in WT. **d)** Difference in amplitude of cycling genes between WT and *cca1lhy*. Outliers are plotted as individual points. Welch’s t-test ****p<0.0001. **e)** Genes that do not meet the cycling threshold in any of the selected circadian datasets in constant light (LL) that are mis-regulated in *cca1lhy* [21,34,53,54]. Clustered using k-means clustering and data is scaled by row. **f)** Expression of *CSU4*, a non-cycling gene from cluster 1 with a genotype effect. **g)** Expression of *PUB56*, a non-cycling gene from cluster 2 with a genotype effect. **h)** Expression of *UBA2A*, a non-cycling gene from cluster 2 with a genotype effect. Gene expression profiles are displayed as average rlog +/- SE. White bars represent subjective day; grey bars represent subjective night. Data shown corresponds to samples collected at 22°C.

More importantly, at the global scale, almost all (98%) genes that are rhythmic in WT exhibit altered rhythms in the mutants (Fig. 1b). The majority of these alterations result in a 12 h shift in time of peak expression between WT and *cca1lhy*, consistent with previous findings of evening phased genes shifting to morning phases in *cca1lhy* (Fig. 1c) [12,51]. Although there are some exceptions to this trend, these results suggest an inversion of clock regulation, where genes that are lowly expressed at a specific time of day in WT may become highly expressed at that time of day in *cca1lhy*, as observed for the other clock genes (Fig. S2a). Of note, this shift in phase is more dramatic than what is observed in diel conditions (∼6h, although more variable throughout the day), highlighting a stronger influence of the clock in constant light conditions and enabling us to identify additional circadian regulation that may be buffered by light cues [52]. In addition to a shifted time-of-peak-expression, genes that cycle in WT have significantly lower amplitudes in *cca1lhy*, implying a less robust clock (Fig. 1d).

To obtain a comprehensive group of cycling genes for downstream systematic analysis, we compared our list of rhythmic genes in WT with published time-resolved transcriptomes under similar circadian conditions (Fig. S2b) [21,34,53,54]. This comparison resulted in ∼1500 genes that are cycling across all datasets (Dataset S1).

### Transcription dynamics of non-cycling genes are controlled by CCA1 and LHY

In addition to regulating cycling genes, CCA1 and LHY have also been shown to bind to the promoters of non-cycling genes; however, the mechanism underlying this regulation remains unknown [12,13]. To better elucidate the genotype effect of mutating *CCA1* and *LHY* outside of their effect on cycling genes, we performed a likelihood ratio test (LRT) comparing the two genotypes and identified ∼5200 genes with a significant genotype effect (Dataset S2). Of these, a subset of 646 genes does not meet the cycling threshold in any of the WT transcriptomes mentioned above but are significantly mis-expressed in *cca1lhy* (Dataset S2). Interestingly, 20% of these are also mis-expressed based on an LRT in a recent diel time-course with *cca1lhy*, suggesting the regulation of these non-cycling genes by CCA1 and LHY is not specific to circadian conditions (Fig. S3a, Dataset S2) [52]. To determine whether these non-cycling and *cca1lhy* regulated genes have similar patterns of expression, we clustered them using k-means clustering to reveal two distinct responses to mis-expression of *CCA1* and *LHY* (Fig. 1e). In cluster 1, genes are more lowly expressed in *cca1lhy* relative to WT, while the reverse is true for cluster 2 (Fig. 1e-h). These two distinct clusters support the dual role of CCA1 and LHY as both activators and repressors. However, the larger proportion of genes in cluster 2 suggests a dominantly repressive role in transcriptional regulation.

While these clusters are not significantly enriched for specific biological processes, genes within these clusters have links to clock components and/or clock-controlled pathways. For example, the transcript abundance of *COP1 SUPPRESSOR 4 (CSU4)* which is a positive regulator of photomorphogenesis and negative regulator of CCA1 is significantly downregulated in *cca1lhy* (Cluster 1, Fig. 1g) [55]. This raises the possibility that CCA1 and LHY activate *CSU4* in a non-rhythmic manner to initiate their own suppression in ambient conditions. In Cluster 2, the expression of three *PLANT U-BOX TYPE E3 UBIQUITIN LIGASES, PUB42*, *PUB55*, and *PUB56*, is significantly increased in *cca1lhy* (Fig. 1h, Fig. S3b-d). None of these three PUBs have previously been linked to clock regulation but they all contain a putative CCA1 binding site in their promoter suggesting that CCA1 may in fact associate to the promoters to regulate their transcription. In support of this, published ChIP-sequencing shows CCA1 occupancy in the *PUB55* promoter [12,13]. Notably, several E3 ligases have been linked to degradation of clock components and also clock regulation; thus, it is possible that the regulation between members of the PUBs and clock genes may involve distinct mechanisms depending on the rhythmicity of these genes [56].

Interestingly, we also identified non-cycling genes within these two clusters with no previous links to the clock that are significantly mis-expressed in *cca1lhy.* These include two of three members of the nuclear RNA-binding protein family *UBP1-ASSOCIATED PROTEIN 2*, *UBA2A* and *UBA2B* (Fig. 1h, Fig. S3d). Given that UBP1 localizes to stress granules, and *UBA2A* forms nuclear foci in response to ABA treatment, this suggests that the clock, mediated by CCA1 and LHY regulation of non-cycling genes, may play a role in mRNA stability or sequestration into stress-induced foci [57].

The notable and significant proportion of non-cycling genes with altered expression in *cca1lhy* raised the question of whether CCA1 and LHY are directly regulating these non-cycling genes through binding to their promoters. First, we found that 4% of these non-cycling genes are targets of CCA1 or LHY based on published ChIP-sequencing datasets (Fig. S3e) [11–13]. An additional 9.5% are targets of other clock components such as the PRRs or TOC1, indicating possible indirect regulation by CCA1 or LHY mediated through other clock genes [58]. Because these ChIP-sequencing experiments represent a single timepoint, some targets may be missed. Next, we screened the promoters of these genes for both EEs and CBSs and found that 30% of these non-cycling genes actually contain a putative CCA1 or LHY binding site (Fig. S3f). These observations highlight a broader and significant role for CCA1 and LHY beyond regulating cycling genes.

To determine whether CCA1 and LHY regulate these non-cycling genes *in planta*, we performed a transactivation assay in *Nicotiana benthamiana*. The promoter of UBA2A was cloned into a dual-luciferase construct and was co-infiltrated with *35S:CCA1-GFP, 35S:LHY-GFP,* or a control construct of *35S:TURBOID-YFP.* We found that co-infiltrating *UBA2A:LUC* with *35S:CCA1-GFP* or *35S:LHY-GFP* resulted in a trend of reduced promoter activity of *UBA2A,* as observed in the transcriptome data, further supporting a role for CCA1 and LHY regulation of non-cycling genes (Fig. S3g).

Recent studies in time-resolved single cell and spatial transcriptomics highlighted that rhythmic expression may be influenced by cell type [59,60]. Given the extent to which non-cycling genes appear to be clock regulated in our above analysis, we considered the possibility that a subset of these non-rhythmic genes may have cell-type specific rhythmicity that is buffered in whole seedlings. To address this, we compared these non-cycling genes with two single-nuclei time course experiments performed in circadian conditions (Fig. S3h) [59,60]. Interestingly, we found that ∼23% (148/646) of the genes that do not cycle in bulk tissue do in fact significantly cycle in one or both single-nuclei datasets (Dataset S2). Of these cell-type specific rhythmic genes, most cycle in either the mesophyll or vascular tissue. A subset of these genes are transcription factors, such as *BHLH35*, which cycles in the epidermis, and has been linked to abiotic stress tolerance [61]. Taken together, this suggests that CCA1 and LHY regulation of non-cycling genes may facilitate cell-type specific rhythms in addition to coordinating constitutive processes.

### A subset of the heat-responsive transcriptome is regulated by *cca1lhy*

Although CCA1 and LHY have been previously linked to the control of heat responsive gene expression in the morning under diel conditions, we questioned if CCA1 and LHY have a functional role in thermotolerance specifically in the morning, as it is the time of their highest expression [14]. We therefore performed an acquired thermotolerance assay on eight-day old Arabidopsis seedlings in the morning (ZT0) and night (ZT12). Seedlings were primed at 37°C for 30 min, followed by recovery at 22°C for 30 min before being subjected to 45°C for 90 min. We found that *cca1lhy* is more susceptible to heat stress compared to WT specifically in the morning (ZT0) (Fig. 2a-b, Fig. S4a-d). Control plants that remained at 22°C or that were only exposed to 37°C for 30 min exhibited no difference in their recovery (Fig. S4b-c). Related to this, a recent study found that *cca1lhy* seedlings have reduced basal thermotolerance only at night (ZT13), further supporting an important role for CCA1 and LHY in coordinating time of day specific thermotolerance in Arabidopsis [62].

**Figure 2:**
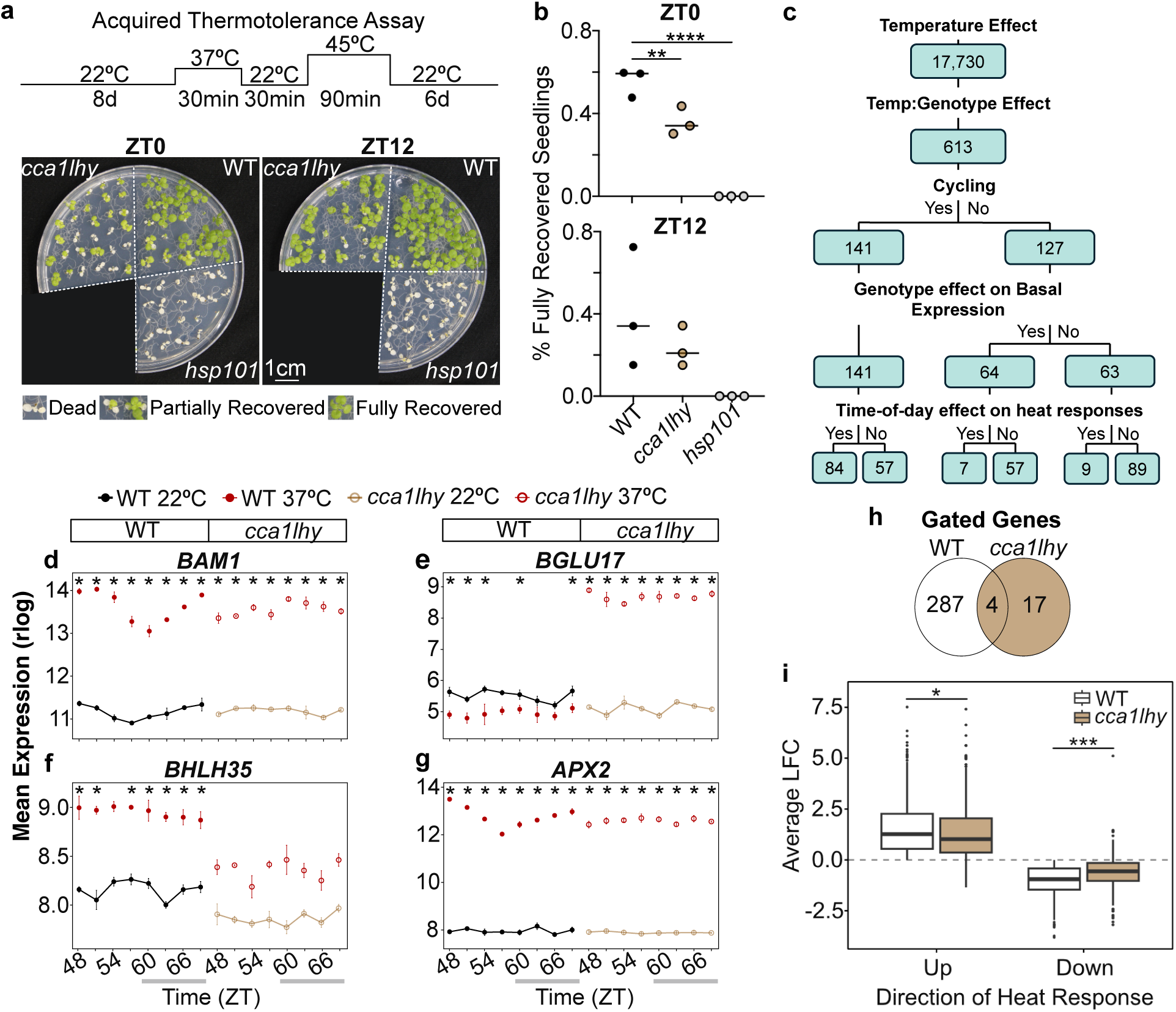
CCA1 and LHY regulate the timing and magnitude of transcriptional heat responses including sulfur-assimilation genes. **a)** Representative images of indicated genotypes (WT, *cca1lhy*, and *Heat Shock Protein 101* mutant, *hsp101*) following acquired heat stress treatment (30 min 37°C, 30 min 22°C, 90 min 45°C) at ZT0 or ZT12. Un-redacted images are provided in Fig. S4. **b)** Quantification of fully recovered seedlings at ZT0 and ZT12. 1-way ANOVA. Dots indicate biological replicates with 3-5 independent plates with ≥15 seedlings per genotype per plate averaged per replicate. **c)** Flow chart displaying significant results of the Likelihood Ratio Test (LRT). Genes with altered temperature responses in *cca1lhy* are separated into cycling and non-cycling groups based on a consensus from 5 time-course transcriptomic datasets [21,34,53,54]. Genotype effect on basal expression derived from the LRT analysis on ambient samples (22°C). Time-of-day effect on heat responses reflects genes with a temperature:time:genotype effect, where time-of-day heat responses differ between genotypes (WT or *cca1lhy*). **d)** Expression profile of *BAM1* **e)** *BGLU17,* **f)** *BHLH35,* and **g)** *APX2*. White and grey boxes represent subjective day and night respectively. Asterix (*) represents significant differential expression (FDR <0.05, LFC >1) at the indicated timepoint. **h)** Gated genes in WT and *cca1lhy*, determined by an LRT within each genotype to determine genes with a temperature:time effect. **i)** Magnitude of heat responses in WT and *cca1lhy*. Average LFC across timepoints was compared. Welch’s unpaired T-test. *p<0.05, **p<0.01, ***p < 0.001, ****p<0.0001.

Given that *cca1lhy* had altered thermotolerance, we next aimed to determine to what extent CCA1 and LHY mediate the gating of heat responsive gene expression throughout the day and under circadian conditions. We expanded our previous time course by collecting WT and *cca1lhy* seedlings after a 1 h 37°C heat treatment, using the same experimental conditions as the ambient samples described above (Fig. S5a). Principal component analysis shows that temperature and genotype explain 63% and 4% of variation between samples respectively (Fig. S1d). Regardless of timepoint, ∼5800 genes were differentially expressed (DEGs) in WT, and ∼5200 in *cca1lhy* (p<0.05, LFC >1) (Dataset S3, Fig. S5b).

To systematically identify genes with altered heat responses in *cca1lhy*, we performed a likelihood ratio test (LRT) to identify the interactions between genotype, temperature, and time (Fig. 2c, Dataset S3). Of the significant heat responsive genes, 613 genes showed a statistically significant genotype effect, suggesting CCA1 and LHY regulate their response to heat stress. The influence of CCA1 and LHY on abiotic stress has largely focused on their regulation of cycling genes [14,63]. However, because of the extent to which CCA1 and LHY regulate non-cycling genes in our above analysis under ambient conditions, we further divided the 613 genes into cycling and non-cycling genes (Fig. 2c). Interestingly, a similar number of genes with altered temperature responses in *cca1lhy* are classified as cycling and non-cycling, emphasizing a role for CCA1 and LHY in regulating heat responses outside of their role in clock-controlled gene expression.

The cycling genes with a genotype effect on their heat responses were mostly enriched for circadian regulated processes (Fig. S5c, Dataset S4). For example, the upregulation and gated response of *BETA-AMYLASE1 (BAM1)* in response to heat stress in WT is significantly altered in *cca1lhy,* particularly in the context of time of day (Fig. 2d). BAM1 has been reported to be involved in transient starch degradation, a pathway controlled by the clock, however, links between BAM1, CCA1, and LHY, and circadian-controlled heat responses were not previously known [64]. Additionally, *BAM1* is targeted by HSFs in ChIP-sequencing data, suggesting both the clock and HSFs may regulate its heat response [65]. These findings suggest that starch degradation and metabolism may be specifically altered in *cca1lhy* under heat stress.

For non-cycling genes with a genotype:temperature effect on their expression, we observe that a subset of genes is also misregulated in *cca1lhy* in ambient conditions (Dataset S3, Fig. 2c). The most striking example of *cca1lhy* mediated alteration of the heat response of a non-cycling gene is *BETA GLUCOSIDASE 17 (BGLU17)* (Fig. 2c and Fig. 2e). In plants, BGLUs are involved in various aspects of stress responses, however the function of *BGLU17* in Arabidopsis remains unknown [66,67]. Here we observed that while the expression of *BGLU17* is modestly reduced in *cca1lhy* compared to WT in ambient conditions, the mRNA abundance is dramatically increased in *cca1lhy* in response to heat stress, while it is slightly reduced in response to heat in WT (Fig. 2e). This clock-specific buffering of heat responses suggests increased *BGLU17* expression may be deleterious to clock function during heat stress.

We also observed that the expression of *BHLH35* is significantly reduced in ambient conditions, and more importantly the upregulation in response to heat stress is significantly dampened in *cca1lhy.* Of note, *BHLH35* was observed to be rhythmic in the epidermis in our above analysis and has been reported to be involved in combined abiotic stress responses [61]. Thus, CCA1 and LHY may play a role in fine tuning the spatial, temporal and temperature response through regulation of *BHLH35*. Clock regulation of *BHLH35* and *BGLU17* has not been previously proposed, emphasizing the importance of considering non-cycling genes as potential targets of circadian control.

To dig further into CCA1 and LHY mediated gating of the heat response, we identified genes that show a time-of-day specific heat response in WT or *cca1lhy*. Strikingly, more than 10 times fewer genes were observed to have a gated heat response in *cca1lhy* compared to WT (Fig. 2h). For example, the gated heat response of *ASCORBATE PEROXIDASE 1B* (*APX2)*, which is known to be involved in ROS homeostasis, is attenuated in *cca1lhy,* most notably between ZT48 - ZT57 (Fig. 2g). In fact, only 4 of the 287 gated genes in WT were also found in *cca1lhy*, suggesting CCA1 and LHY are required for the majority of time-of-day regulation of heat responses regardless of the time of day when the gating occurs (Fig. 2h).

Furthermore, in addition to influencing heat responses throughout the day, we observed that regardless of whether genes were upregulated or downregulated, CCA1 and LHY contributed to the intensity or magnitude of the heat responses, where genes are generally less up- or down-regulated by heat in *cca1lhy* (Fig. 2i). Together, this emphasizes that CCA1 and LHY are important regulators of the timing and intensity of heat responses for a subset of genes.

We hypothesize that CCA1 and LHY mediated regulation of the timing and intensity of heat stress responses may be due to the binding of CCA1 and LHY to target promoters. Integrating published ChIP-sequencing data revealed that CCA1 and LHY occupy the promoter of 17% of the genes with a genotype:temperature effect on their expression, with the caveat that CCA1 and LHY binding may be dynamic or altered under heat stress conditions (Fig. S6a) [11–13]. A systematic analysis revealed that of genes with altered heat responses in *cca1lhy*, 36% have a putative CCA1 or LHY binding site (EE or CBS) in their promoter (Fig. S6b). Cycling genes with altered heat responses are slightly more likely to contain an EE in their promoter than non-cycling genes with altered heat responses (43% vs 30%). We also found that 21% of genes with altered heat responses contain a G-Box motif (CACGTG) in their promoters, suggesting that regulation through other clock components such as the PRRs or other TFs may contribute to the observed responses (Fig. S6b). Together, these results support that the altered heat response for a subset of genes in *cca1lhy* may result from loss of CCA1 and LHY binding to their promoters. However, given that CCA1 and LHY ChIP-sequencing targets have varied levels of perturbed heat responses in *cca1lhy*, CCA1 and LHY likely also influence other heat-responsive regulatory pathways (Fig. S6c).

### Specific and combined circadian coordination of heat responsive pathways

Clock coordination of heat stress responses is likely to include multiple mechanisms with specificity for key hubs depending on the intensity and duration of the heat stress. Based on previous studies it is known that heat stress responses may be mediated by Heat Shock Transcription Factor (HSF)-dependent or -independent pathways and clock components have been linked to these alternative regulatory mechanisms [50]. Given this, we next explored whether CCA1 and LHY modulation of heat-responsive gene expression is independent of or through known thermoregulators. We found that ∼59% of the genes with altered heat responses in *cca1lhy* contain at least one Heat Shock Element (HSE) in their promoter, suggesting they may be bound by an HSF. Both *HSFA3* and *HSFA8*, have altered ambient expression in *cca1lhy*, while *HSFA1E* has altered heat responses (Fig. 3a, Fig. S7a-b). Additionally, other HSFs are differentially expressed depending on time of day, suggesting that HSF-mediated heat responses may be altered in *cca1lhy*.

**Figure 3:**
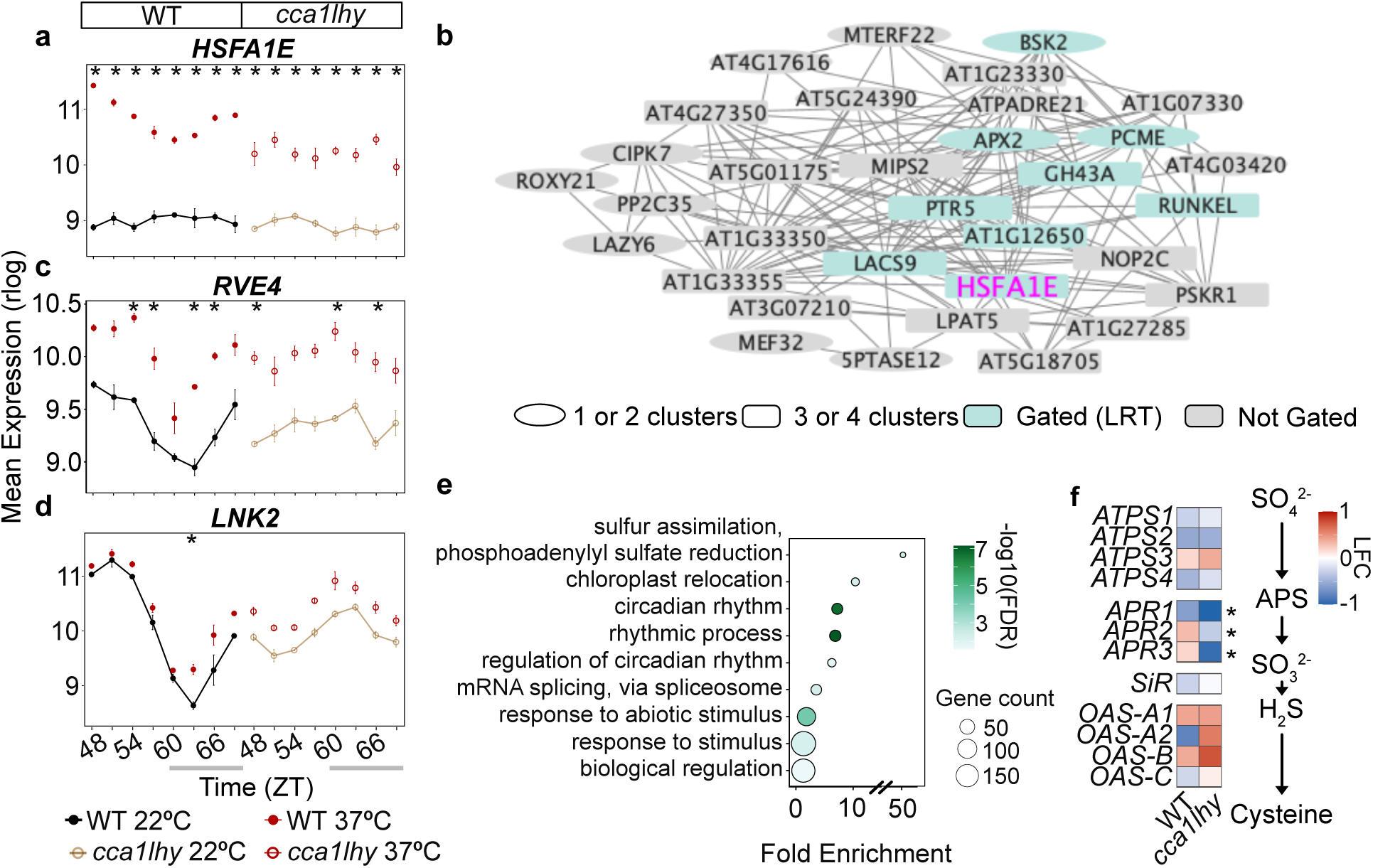
CCA1 and LHY influence gating of heat responses through HSF-dependent and - independent pathways. **a)** Expression profile of *HSFA1E*. **b)** Genes co-expressed with HSFA1E. Blue indicates a gene with altered gating between WT and *cca1lhy*. Ellipse = in one or two co-expression modules containing *HSFA1E*, rectangle = in three or four modules. Nodes arranged manually for visibility. **c)** Expression profile of *RVE4* and **d)** *LNK2.* White and grey boxes represent subjective day and night respectively. Asterix (*) represents DEG (FDR <0.05, LFC >1) at the indicated timepoint. **e)** GO term enrichment of the 613 genes with altered temperature responses in *cca1lhy*. **f)** Heatmap of genes related to sulfur assimilation in Arabidopsis [128]. * Indicates a significantly different temperature response in *cca1lhy* based on the LRT. Data is shown for ZT48. *ATPS = ATP SULFURYLASE*, *APR = APS REDUCTASE*, APS = adenosine 5’-phosphosulfate, *SiR = SULFITE REDUCTASE, OAS = O-ACETYLSERINE (THIOL) LYASE*.

HSFA1E has been previously shown to act redundantly with other HSFAs, such as HSFA1D in the activation of *HSFA2* [68]. To identify if genes with altered heat responses in *cca1lhy* may act in a HSF-dependent manner mediated by HSFA1E, we generated coexpression modules using ClustONE. We found that *HSFA1E* is coexpressed with several genes with gated heat responses that differ between WT and *cca1lhy* where the highest heat response occurs in the morning in WT (Fig. 3b, Dataset S3). For example, *ASCORBATE PEROXIDASE 2 (APX2)* a non-cycling gene, is important for ROS scavenging under heat stress, and *MYO-INOSITOL-1-PHOSPHATE SYNTHASE* 2 (*MIPS2)* which produces *myo*-inositol and has several functions in abiotic stress tolerance (Fig. 3b). The coexpression of these genes with *HSFA1E* suggests that HSFA1E may contribute to their regulation or that both *HSFA1E* and its co-expressed genes share common upstream regulators, which may include CCA1 or LHY.

It is also possible that CCA1 and LHY mediate heat stress gating through non-HSF-dependent pathways involving other daytime expressed clock proteins such as RVE4 and RVE8, as well as their transcriptional coactivators LNK1/LNK2 [50,62]. All four of these clock components have altered expression in *cca1lhy* specifically under ambient conditions (Fig. 3c and 3d, Fig. S8a-d). The RVEs have been shown to be important for the induction of *ERF DOMAIN 53 (ERF53)* and *ERF54* in response to heat during the day, affecting seedling survival [50]. We observed that *ERF53* and *ERF54* have altered patterns of heat response in *cca1lhy*, with *ERF53* generally less heat responsive in *cca1lhy* compared to WT (Fig. S8c-d). Similarly, overexpression of LNK1 and LNK2 have been shown to increase the abundance of *HSFA8 [62]*. We speculate that since *LNK1* and *LNK2* transcript abundance is lower at ZT48 in *cca1lhy*, and *HSFA8* is more highly expressed, CCA1 and LHY may regulate the expression of *HSFA8* independent of the LNKs (Fig. S8e).

Regardless of whether gated heat responses are directly or indirectly regulated by CCA1 and LHY, we found that genes with altered heat responses in *cca1lhy* are involved in sulfur assimilation, chloroplast localization, circadian processes, mRNA splicing via spliceosome, and response to stimulus (Fig. 3e, Dataset S4). While links between clock components or circadian regulation of most of these processes has been previously reported, the impact of an altered clock and heat stress is mostly unexplored [21,34].

Notably, changes in the sulfur assimilation pathway is associated with circadian control of temperature responses, and sulfur-containing compounds have well established roles in ROS scavenging in response to stress [14,21,69,70]. The three-member family of ADENOSINE 5’-PHOSPHOSULFATE REDUCTASES (APRs) represent the rate limiting step in the sulfur assimilation pathway in Arabidopsis [65,70]. Interestingly, we observed that all three members, *APR1*, *APR2* and *APR3,* are significantly downregulated in *cca1lhy* after heat treatment relative to WT at ZT48 (Fig. 3f). This suggests that altered sulfur metabolism in response to heat stress in *cca1lhy* may alter ROS scavenging and compromise thermotolerance. Altered regulation of transcriptional heat responses, dependent or independent of known regulators, likely results in gene expression dynamics that persist through multiple levels of regulation to impact overall thermotolerance.

### Mis-expression of CCA1 and LHY alters the heat responsive proteome

In a previous study, we found that the clock gates the heat response of a significant proportion of genes at the translation level in Arabidopsis [21]. Given that we observe that genes involved in RNA splicing differ in response to heat stress in *cca1lhy* and previous work suggesting CCA1 may regulate ribosome loading, we next investigated to what extent CCA1 and LHY mediated gating of heat responses impacts the proteome (Fig. 3e) [36]. Briefly, WT and *cca1lhy* seedlings were grown in similar light and temperature conditions as the transcriptome analysis described above. However, samples were collected following heat treatment at the two times of the day where *CCA1* and *LHY* are normally at their peak or trough expression (ZT48 and ZT60 respectively) and Liquid Chromatography-Tandem Mass Spectrometry (LC-MS/MS) was performed (Fig. 4a).

**Figure 4:**
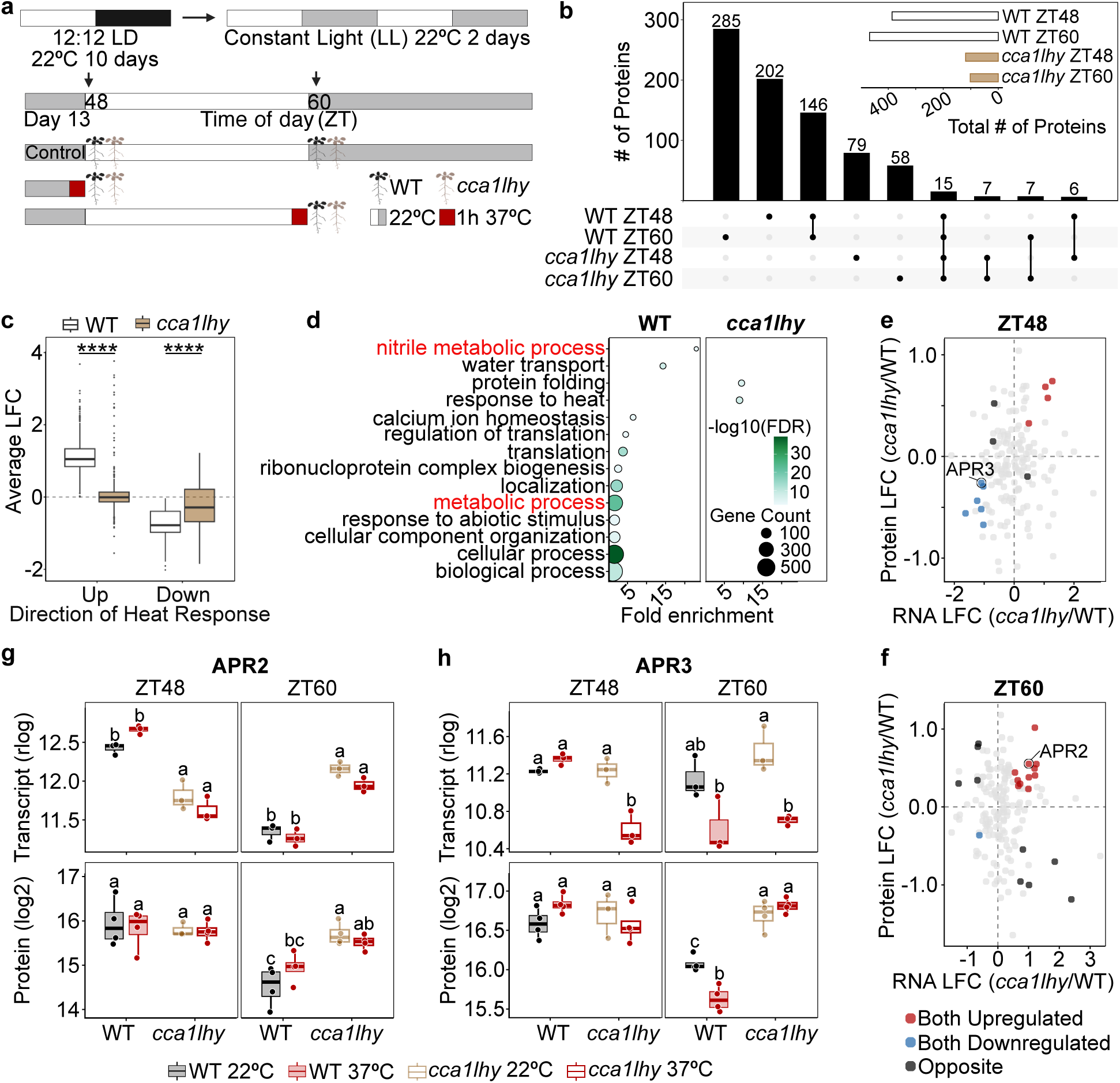
Heat responses altered in the proteome reflect transcriptional and post-transcriptional regulation. **a)** Experimental design. 1 h 37°C heat stress performed ending at ZT48 and ZT60. **b)** Differentially abundant proteins (DAPs) within each genotype and timepoint. Select intersects shown. **c)** LFC of heat responses in WT and *cca1lhy* of all DAPs in either genotype (WT or *cca1lhy*). Data is averaged across both timepoints. Welch’s unpaired T-test. ****p < 0.0001. **d)** GO enrichment of DAPs in WT and *cca1lhy*. Circle size indicates gene number, color indicates - log10(FDR). **e)** Differential expression of *cca1lhy* compared to WT in both the transcriptome and proteome at ZT48 and **f)** at ZT60. LFC from pairwise comparisons comparing the heat-treated samples between the two genotypes. Dots are colored if the FDR (transcriptome) and q (proteome) is <0.05. Black circles indicate significance in both the transcriptome and proteome with opposite directions of LFC. Genes are displayed if they have a genotype effect on their temperature response based on the LRT in the RNA-seq time-course. **g)** Transcript and protein abundance of APR2 and **h)** APR3 at Z48 and ZT60. Significant differences based on a 2-way ANOVA with Tukey’s significant difference test indicated by lowercase letters (p<0.05).

In ambient conditions, of the 6729 proteins quantified, 15% differ in abundance between WT and *cca1lhy* at ZT48 (995 proteins) and ZT60 (982 proteins) (q<0.05, LFC>1) (Fig. S9a-b, Dataset S5). Some of these differentially abundant proteins (DAPs) may reflect cycling proteins with altered phase in *cca1lhy,* such as MIPS1, which was found to be rhythmic in a proteome in circadian conditions [37]. Because a time-resolved proteomic dataset in circadian conditions with sufficient resolution to identify rhythmic proteins does not exist, this limits accurate determination of cycling proteins. Yet, fewer DAPs vary within *cca1lhy* between ZT48 and ZT60 compared to WT, suggesting that cycling is being perturbed in the proteome (Fig S9).

To distinguish changes that are largely driven by regulation in the transcriptome, we compared both transcript and protein abundance differences between *cca1lhy* and WT at each timepoint. Based on this, we identified ∼300 genes that showed a similar increase or decrease in mRNA and protein abundance in *cca1lhy* compared to WT suggesting that the genotype-specific transcriptional regulation persists through the proteome (Dataset S5, Fig. S9c-d). Interestingly, of the 300 concordant genes, we identified 11 genes that show similar differences in expression between the genotypes in the transcriptome and proteome at both timepoints (Dataset S5). Notably, UBA2A and UBA2C, which we discussed above, are upregulated in *cca1lhy* in both the transcriptome and proteome. Given that *UBA2A* expression is non rhythmic in the transcriptome, this suggests that the regulation of non-cycling genes by CCA1 and LHY may persist through the proteome (Fig. S9f). We also found that CSU4, another non-cycling gene discussed above with altered expression in *cca1lhy*, is similarly downregulated in both the transcriptome and proteome at night (Fig. S9g). However, in the morning the protein abundance of CSU4 is not significantly altered in *cca1lhy*, suggesting that this gene is regulated at additional levels after transcription.

In comparison, most of the concordant genes are similarly mis-regulated in *cca1lhy* under diel conditions (at ZT1 and ZT13) in the transcriptome, but not in the proteome based on published data (Fig. S9e) [52]. This may suggest that the timing between mRNA production and protein synthesis differs in diel and circadian conditions or that light signaling produces cycles that persist in the *cca1lhy* proteome in diel conditions.

In response to heat stress, 386 and 468 proteins are heat responsive in WT at ZT48 and ZT60, respectively, while 119 and 102 proteins were responsive in *cca1lhy* (Fig. 4b, Dataset S5). By comparing only within genotypes, we minimize conflating a phase change of protein expression or heat response with a genotype effect. Interestingly, the lower number of differentially abundant proteins (DAPs) in *cca1lhy* suggests that CCA1 and LHY may regulate the magnitude of heat responses similar to our observations in the transcriptome. Thus, to comprehensively assess the relevance of CCA1 and LHY on the proteome heat response, we compared the LFC of all DAPs in either WT or *cca1lhy*. Strikingly, we found that proteins upregulated in WT on average became less upregulated in *cca1lhy*, while proteins downregulated in WT were less downregulated in *cca1lhy* (Fig. 4c). DAPs in *cca1lhy* are specifically enriched for heat responses, while multiple processes are enriched in WT, including metabolic processes and nitrile metabolic processes (Fig. 4d, Dataset S4). Together these observed modulations of heat responses in the *cca1lhy* proteome emphasize the importance of circadian regulation across multiple levels of regulation.

We then identified genes whose expression is altered in *cca1lhy* relative to WT that show similar or distinct mRNA and proteome abundance changes (up or down regulation in *cca1lhy* vs WT) in response to heat stress (Fig. 4e-f, Dataset S5). About 20 genes have altered heat-responses in *cca1lhy* compared to WT that are significant in both the transcriptome and proteome. However, even within these genes that are both a DEG in the transcriptome and DAP in the proteome (*cca1lhy* 37°C vs WT 37°C, FDR or q <0.05), we observed indications of post-transcriptional regulation. For example, two of the previously discussed APRs, APR2 and APR3, have similar patterns of expression in the transcriptome and translatome at ZT60 and ZT48 respectively (Fig. 4g-h). For APR2, differences between the genotypes are similar in the transcriptome and proteome at ZT60, however genotype differences in the transcriptome at ZT48 are lost in the proteome. Similarly, APR3 mRNA and protein abundance is lower in *cca1lhy* relative to WT after heat stress at ZT48. Of significance, much of the intra-genotype response to heat stress is attenuated in the *cca1lhy* proteome at both timepoints. These examples highlight both that *CCA1* and *LHY* dependent heat-responsive transcriptional regulation may persist in the proteome, and that additional post-transcriptional regulation is likely occurring.

Genes that behave differently between the transcriptome and proteome suggest that additional layers of regulation may be influenced by CCA1 and LHY in response to heat stress. To further explore this, we leveraged a Translating Ribosome Affinity Purification sequencing (TRAP) dataset generated under similar experimental conditions as the proteomics analysis using a *cca1-1 lhy-20 35S:HF-RPL18* line (*cl* TRAP), where the *cca1* allele is segregating resulting in a knock-down of *cca1* (Fig. 4a, Fig. S10a, Dataset S6). We clustered genes based on their behavior in the transcriptome, translatome, and proteome within each genotype. This comparison revealed that most genes seem to behave similarly in the transcriptome and translatome but differ in their heat-stress proteome response (Fig. S10c). The similarity between the transcriptome and translatome may result from the TRAP line being only a partial knock-down of CCA1, however the proteome still responds strikingly differently compared to the transcriptome. For example, *TONOPLAST INTRINSIC PROTEIN 2* (*TIP2*), as well as other aquaporins in the *TIP* and *PLASMA MEMBRANE INTRINSIC PROTEIN (PIP)* families are highly induced by heat in both the transcriptome and translatome regardless of genotype (Fig. S10d). However, in the proteome, the heat induction of TIP2 protein abundance is significantly lost in *cca1lhy*, indicating possible differences in the time between ribosome binding and protein production or in post-translational regulation (Fig. S10d). Post translational regulation of multiple aquaporins, including several members of the PIPs, is known [71,72]. For example, PIP2;1 is reported to undergo phosphorylation and ubiquitination in response to osmotic stress, altering its localization and stability [73,74]. The observed cross layer dynamics for this subset of genes implicates new connections between the clock and post-translational regulation that may be mediated by CCA1 and LHY at high temperatures. Taken together, by comparing heat responses across multiple layers of gene regulation we observed that both the magnitude and direction of heat responsive gene expression is notably more disrupted in the proteome compared to the transcriptome in *cca1lhy*.

### Influence of CCA1 and LHY on the dynamics of primary metabolites in response to heat stress

Both the circadian clock and abiotic stress are known to alter metabolic processes [75–80]. In response to heat, we observed aspects of primary metabolism including sulfur assimilation and starch degradation altered in *cca1lhy* across different levels of regulation, as well as different responses of proteins involved in metabolic processes. We therefore performed targeted metabolomics on central carbon metabolites to reveal compounds altered in *cca1lhy* and WT following a short (1 h ending at ZT48) or extended (3 h ending at ZT50) 37°C heat stress in the morning (Fig. 5a, Dataset S7).

**Figure 5:**
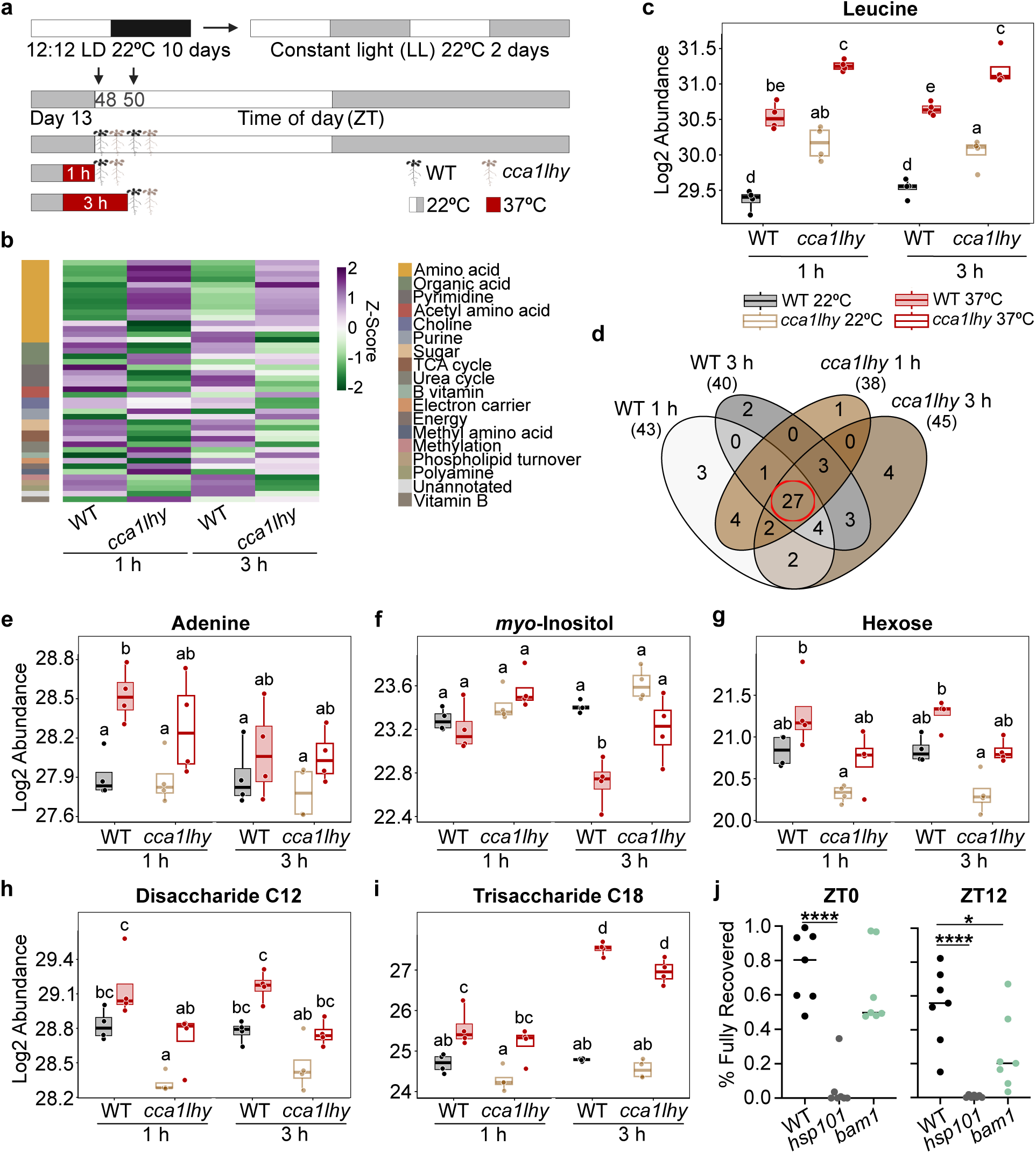
Altered metabolites in *cca1lhy* may reflect differences in starch degradation. **a)** Experimental design. 1 h and 3 h heat treatments (37°C) performed and ended at ZT48 or ZT50. **b)** Heatmap of metabolites that significantly differ in pairwise comparisons within or between genotypes. Only ambient conditions (22°C) are shown. Scale shows z-score of log2 normalized relative abundance. **c)** Relative abundance of leucine. Each dot represents a biological replicate (N=4). Significant differences based on a 3-way ANOVA with Tukey’s significant difference test indicated by lowercase letters (p<0.05). **d)** Overlap between heat-responsive metabolites in WT and *cca1lhy*. **e)** Relative abundance of adenine, **f)** *myo*-inositol, **g)** hexose, **h)** disaccharide C12, and **i)** trisaccharide C18. **j)** Quantification of fully recovered seedlings in thermotolerance assays at ZT0 and ZT12. Genotypes were WT, *bam1*, and *Heat Shock Protein 101* mutant (*hsp101*). Dots represent biological replicates (N=7) with ≥3 plates per treatment within each replicate with ≥15 seedlings per genotype per plate. One way ANOVA ***p<0.001 ****p<0.0001. Representative images in Fig. S12.

We found that genotype explained 22% of variation between the samples, suggesting *cca1lhy* has a distinct metabolic profile compared to WT (Fig. S11a). Overall, we identified ∼35% of the detected metabolites differ between *cca1lhy* and WT at ZT48 in ambient conditions and 22% at ZT2 (Fig. 5b, Dataset S7). Notably, several of these metabolites are amino acids that have increased abundance in *cca1lhy*, such as leucine (Fig. 5c). We speculate that this may be due to premature degradation of starch, which has been reported in *cca1lhy*, that results in protein catabolism leading to increased levels of amino acids [76,81,82]. We also found that sugars including hexoses and disaccharides C12 are less abundant in *cca1lhy* in ambient conditions, reflecting reduced energy availability. Altered abundances of these metabolites may also reflect changes in rhythmicity, as several amino acids such as lysine and valine have been shown to cycle in free-running conditions [77].

In response to heat stress, ∼40 metabolites have altered abundance in each genotype at each timepoint (Fig. 5d). Most metabolites are responsive within both genotypes under both temperature treatments. Interestingly, ten metabolites are specifically responsive to heat in *cca1lhy* or WT, including adenine, which is specifically upregulated in WT (Fig. 5e, Dataset S7). Increased adenine abundance has been linked to higher thermotolerance, and so reduced induction in *cca1lhy* may contribute to reduced recovery at ZT0 [83].

Of note, some metabolites are also specifically altered after a longer duration of heat stress. For example, *myo*-inositol is less abundant only after 3 h of heat treatment (Fig. 5f). *Myo-*inositol is also generally more abundant in *cca1lhy* in all conditions, leading to increased abundance relative to WT after heat stress. We had also observed that *MIPS2*, a *myo-*inositol biosynthetic gene, is co-expressed with *HSFA1E*, indicating circadian control of its synthesis may occur in an HSF-dependent manner. Additionally, several proteins involved in inositol-related metabolism are differentially abundant between WT and *cca1lhy* in our proteomic analysis, including MIPS1 and four proteins involved in phosphatidylinositol synthesis, suggesting that cell membrane remodeling may be altered in *cca1lhy*.

Interestingly, sugar metabolism remains altered after heat stress in *cca1lhy*. For example, the abundance of hexose, disaccharide C12, and trisaccharide C18 sugars are reduced in *cca1lhy* relative to WT, although are all still induced by heat (Fig. 5g-h). Some sugars, such as the trisaccharide raffinose have been demonstrated to increase in response to heat stress, and possibly function as osmolytes, ROS scavengers, or serve to re-allocate carbon [84]. We therefore hypothesize that altered sugar abundance in *cca1lhy* may contribute to reduced thermotolerance.

Given the altered carbon metabolism observed in the analysis above, we further examined alterations related to the starch degradation pathway, as usable sugars at dawn are derived from starch degradation as opposed to novel synthesis. We found that of putative starch degrading enzymes, *BAM1*, discussed previously, is dramatically induced by heat and is also mis-expressed in *cca1lhy* in both the transcriptome and proteome (Fig. S12a-b). Using a *bam1* mutant line, we performed a similar thermotolerance assay as described above in Fig. 2. We found that *bam1* seedlings are indeed more susceptible to heat treatment than WT (Fig. 5j and S12c-d). As *bam1* abundance is reduced in *cca1lhy* at ZT48, we speculate that this may contribute to the reduced recovery observed in *cca1lhy*. This result suggests that *cca1lhy* may regulate thermotolerance by controlling energy availability through the day.

## Discussion

Optimal responses to high temperature rely on a robust and properly functioning circadian clock. This complex regulatory relationship spans across most if not all layers of gene regulation and is likely to involve multiple mechanisms throughout the day. In this study, we generated and combined transcriptomics, proteomics, and metabolomics to determine specific and shared regulation and responses mediated by the core clock components CCA1 and LHY and other known heat stress related clock genes involved in thermotolerance (Fig. 6a-b).

**Figure 6:**
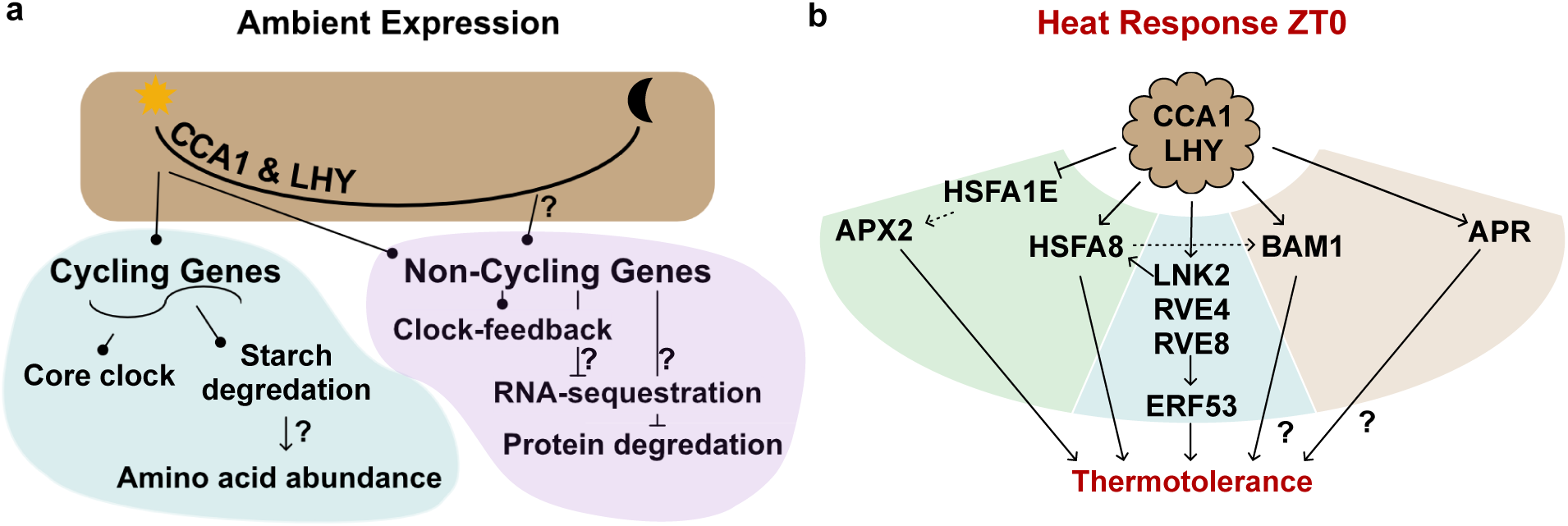
CCA1 and LHY regulate diverse processes in ambient (22°C) and heat stress (37°C) conditions. **a)** Regulation and pathways altered or influenced by CCA1 and LHY in ambient conditions or **b)** in response to heat stress. Question mark (?) indicates suggested but unproven regulation. Background colors in b) reflect CCA1 and LHY regulation acting through the Heat Shock Transcription Factors (HSFs), REVIELLEs (RVEs), or potential independent pathways. Arrow indicates activation, bar indicates repression, and circle indicates varied or unclear regulation.

In ambient conditions, we observed that CCA1 and LHY regulate both cycling and non-cycling genes, which reinforces and further expands our knowledge of what genes may be considered clock-regulated (Fig. 1 and Fig. 6a). CCA1 binding to arrhythmic genes has been previously observed, and in fact, clock proteins binding to non-cycling genes is also common in other biological systems. For example, in the mouse liver, 40% of BMAL1 bound genes are not significantly rhythmic [13,85]. This may result from differences in mRNA stability, where stable transcripts buffer the input of rhythmic transcription, which can be investigated using nascent RNA-seq [86]. It is also possible that this may result from cell-type specific rhythmicity [13,59]. Our findings support this hypothesis as 23% of the non-cycling genes misregulated in *cca1lhy* have tissue-specific rhythms (Fig. S3g). This raises the possibility that CCA1 and LHY function extends to controlling cell-type specific targets as has been seen for clock transcription factors in other organisms, which may be resolved by performing a time-resolved tissue-specific ChIP-sequencing experiment [87,88]. Tissue-specific CCA1 and LHY function would support the presence of a spatially heterogeneous clock in Arabidopsis where distinct cell types have different phases of expression [89]. Heterogeneity in heat responses in root tissue has also been identified, which may be fine-tuned by differences in the circadian state in different cells [90].

We demonstrate that a subset of heat-responsive transcripts has altered responses to heat in *cca1lhy* (Fig. 2a). Some of these genes are also shared with our previous data in diel conditions, although to a limited extent, which is possibly due to differing light conditions and timepoints [14]. Interestingly, transcripts with altered heat responses in *cca1lhy* are almost evenly divided between cycling and non-cycling genes, suggesting a role for CCA1 and LHY in regulating heat responses outside of their established clock functions (Fig. 1a). Although CCA1 and LHY are known to regulate stress responses, such as ABA biosynthesis and signaling, it has been previously unclear how they regulate heat responses [11,14,63]. Our findings suggest that CCA1 and LHY likely regulate transcriptional heat responses partially through the regulation of both the ambient and heat-responsive expression of HSFs, especially HSFA8 and HSFA1E (Fig. S5, Fig. S6) [14,62]. CCA1 and LHY are similarly likely to regulate HSF-independent regulation, such as that driven by RVE4 and RVE8, as they are mis-expressed in our above analysis (Fig. 3, Fig. S7). Within the heat-responsive transcriptome, a proportion of genes have gated heat responses based on the time-of-day, which is almost entirely lost in *cca1lhy* (Fig. 2h). While most of these transcripts are rhythmic, we observed no enrichment for their phase (timing of peak expression) in ambient conditions, which is consistent with previously observed patterns of circadian gating [21]. Non-cycling genes also lost gating in *cca1lhy,* especially genes that were less heat responsive at the end of day, suggesting CCA1 and LHY promote their expression in response to heat. The loss of gating in *cca1lhy* regardless of the phase suggests that, perhaps unlike the RVEs, which act at a specific time of day, that CCA1 and LHY may have a broader regulatory role in controlling circadian gating [50]. CCA1 and LHY may also associate with other proteins to affect heat responses, potentially in a time-specific manner. It has been recently suggested, for example, that the clock protein LIGHT-REGULATED WD 1 (LWD1) forms distinct protein complexes throughout the day, while the RVEs interact with distinct proteins to regulate cold and heat-responses [46,91,92]. It would therefore be interesting to identify CCA1 and LHY protein interactors throughout the day and in response to stresses to identify additional proteins involved in regulating heat-responsive gating.

We also identify alterations in the heat-responsive proteome, where the proteomic response to heat stress seems to be highly time-point specific, with time-of-day differences in heat responses persisting in *cca1lhy* (Fig. 4b). This suggests that gating may be more strongly regulated in the proteome than in the transcriptome or also may reflect changes in ambient expression that do not occur, or occur at a different phase, between the transcriptome and the proteome [21,52]. As the translatome seems globally similar to the transcriptome when *CCA1* and *LHY* expression is perturbed, circadian regulated protein stability and degradation may differentiate heat responses in the *cca1lhy* proteome. Future experiments using proteosome mutants may help to clarify if the difference in *cca1lhy* protein responses to high temperature are dependent on differential protein degradation.

Interestingly, although we observe *cca1lhy* specific regulation in the transcriptome and the proteome, there is little overlap between regulation in the two datasets. While this is not unexpected as previous studies have also found low correlation based on the numerous levels of regulation between mRNA and protein abundance, it suggests that gating in the transcriptome and proteome may have different consequences [93]. Circadian gating of transcription in response to other stress, such as of the *CBFs* in response to cold and of *ABF3* during drought stress, has effects on stress tolerance [41,63]. In addition, Arabidopsis plants tolerate heat stress differently at different times of day suggesting that gating, including transcriptional gating by RVE4 and RVE8, has a functional role in thermotolerance [42,50]. Therefore, transcriptional gating is important for stress tolerance and our time-resolved analysis and results demonstrates a role for CCA1 and LHY in mediating this process. However, at the physiological level, how this partitioning results in time-restricted thermotolerance requires more whole and single cell genome-scale data and future functional studies.

Our comparative -omics analysis revealed processes that are consistently mis-regulated in the transcriptome and proteome in *cca1lhy* in response to heat such as sulfur assimilation and starch degradation (Fig. 3f, Fig. S12). Altered sulfur assimilation may influence the abundance of sulfur-containing molecules, including glutathione, which is involved in ROS scavenging [70]. It may also represent crosstalk with nitrogen assimilation, which is moderated by ELONGATED HYPOCOTYL 5 (HY5), both of which are known to be clock-regulated and have altered heat responses in our data [94]. Although we did not observe altered glutathione levels in our metabolomics analysis, we hypothesize that this may be due to the collection time limiting quantification of newly synthesized metabolites and instead reveal fluxes between metabolites that are largely unaltered in *cca1lhy*. Similarly, altered starch degradation in response to heat in *cca1lhy* may correspond to altered thermotolerance. These pathways, as well as other non-cycling candidates mentioned above, *UBAs, CSU4s, BHLH35,* and *PUBs*, should be considered as candidates for future functional characterization of their interaction with the circadian clock. Additionally, these targets should be considered as inputs for modelling efforts to better understand clock function and outputs [95,96].

## Conclusions

This comprehensive and integrated data revealed common and distinct patterns of heat response in the transcriptome, proteome, and metabolome throughout the day. In addition, new target genes, proteins, and metabolites linking the clock and heat responses that may contribute to time-specific thermotolerance were identified. Future work that includes targeted genetic manipulation to alter circadian regulation of multiple heat stress responsive pathways may enable engineering of plants that are more tolerant to nighttime or extreme high temperatures.

## Methods

### Plant Materials and Growth Conditions

In all experiments, *Arabidopsis thaliana* Columbia-0 (Col-0) was used as wild-type (WT) unless noted otherwise. The *cca1-1 lhy-20* line (*cca1lhy*), *cca1-1 lhy-20 35S:HF-TRAP, and 35S:HF-TRAP* have been previously reported [14,21,97]. A T-DNA insertion in *BAM1* (Salk_039895) was ordered from the Arabidopsis Biological Resource Center, was confirmed to be homozygous and knocked-out using PCR and RT-qPCR, respectively (Table S3). Unless otherwise stated, seeds were surface sterilized and stratified at 4°C for two to three days in the dark. Seedlings were grown in circular petri dishes on filter paper placed on Murashige and Skoog (MS) medium supplemented with 1% (w/v) sucrose. Plates were randomly arranged in the growth chamber, and seedlings were grown for 10 days in 12 h white light (F17T8/TL841 Fluorescent Tube, ∼130 μEm-2s-1) and 12 h dark (LD) cycles (12:12 LD) at constant 22°C temperature. On day 11, plants were transferred to constant light (LL) for two days.

#### Ambient and Heat stress treatment

For the RNA-sequencing time course, a 1 h heat treatment at 37°C were performed on different sets of WT and *cca1hy* seedlings every 3 h (ZT48-ZT69) starting at the time of lights ON for day 13 [98]. For clarity, all heat treatments for the -omics experiments were performed to end at the ZT indicated, for example a heat-treated sample labeled ZT51 was heat treated beginning at ZT50 and collected at ZT51. Ambient and heat-stressed samples were collected immediately after the 1 h heat stress. For the TRAP and proteomics experiments 1 h 37°C heat treatments were performed ending at ZT48 and ZT60 on day 13 before immediate collection. For the targeted metabolomics experiments, WT and *cca1lhy* tissue was collected at ZT48 (1 h 37°C heat treatment) and at ZT50 (3 h 37°C heat treatment). Three biological replicates were grown for each condition and time point for RNA-seq. Four replicates were used for proteomic and metabolomic experiments. Replicates correspond to different plates, randomly arranged in the growth chamber on different shelves. Control or stressed seedlings were snap frozen in liquid N2 and stored at -80°C until sample processing.

### Heat Stress Thermotolerance Assays

Seeds were surface sterilized and plated on circular petri dishes (100x15mm) containing 25 ml of Murashige and Skoog (MS) and lacking additional sucrose. After cold-stratification at 4°C for 2 - 3 days, plants were grown in 12:12 LD conditions at 22°C. On the 8th day of growth, heat treatments were performed. The heat treatment was 30 min 37°C, 30 min 22°C, 90 min 45°C and was performed in a growth chamber with control plates kept at 22°C or exposed to only the 30 min 37°C heat treatment. All heat treatments were performed in the light, with all plates involved in the experiment returned to normal light:dark cycles immediately upon completion of the 45°C stress. Plants were allowed to recover for 6 days before assessment of recovery was performed. Recovery was graded as “dead” or the seedling was fully bleached, “partially recovered” where at least the new leaves and apical meristem remained green, and “fully recovered” where no bleaching or damage was observed.

### RNA Extractions, Library Preparation and Sequencing

Total RNA was extracted using the GeneJET Plant RNA Purification Mini Kit (Thermo Fisher Scientific). The resulting RNA was DNase I treated to remove DNA contamination (Thermo Fisher Scientific) and 1 ug of total RNA was mRNA purified using dynabead oligoDTs [99]. RNA libraries were prepared as previously described using modifications previously detailed [14,100]. A final amplification step of 14 cycles was performed using index adapter enrichment primers. Final libraries were purified using 0.9 volume Ampure XP beads. Final libraries were then quantified using a Qubit 2.0 Fluorescence Reader (Thermo Fisher Scientific) and quality and quantification was confirmed using an Agilent TapeStation 4150. Resulting libraries were multiplexed and sent for 150 bp paired-end illumina sequencing on a NovaSeq XPlus 10B lane (Novogene).

### Translating Ribosome Affinity Purification, mRNA isolation, Library Preparation and Sequencing

TRAP was performed as previously described [21,101]. Briefly, working in a cold-room, 1 volume (1-2 mL) of powdered tissue was homogenized in 5 volumes of Polysome Extraction Buffer (0.2M Tris-HCl pH9, 0.2M KCl, 0.025M EGTA pH8, 0.035M MgCl_2_, 1 mM DTT, 1 mM PMSF, 100 ug/mL cycloheximide, 50ug/ml chloramphenicol, 1% PTE (v/v), 0.2% (w/v) polyoxyethylene(23)lauryl ether, 0.2% (v/v) Triton X-100), 0.2% (v/v) octylphenyl-polyehtylene glycol, and 0.2% (v/v) polyoxyethylene sorbitan monolaurate 20) using a glass homogenizer. The homogenized sample was transferred to a cooled polycarbonate tube before centrifugation at 16,000 x g for 15 min at 4°C. The supernatant was then filtered through a layer of miracloth to obtain the clarified extract. The total fraction of RNAs was later extracted from a reserved 200 ul aliquot of the clarified extract. The remaining extract was co-incubated with a-FLAG M2 (Millipore-Sigma) coupled Protein G Dynabeads (Thermofisher) for 2 h at 4°C with gentle rocking. Beads were then separated using a magnet for 4 min at 4°C and were washed using 6 mL of Wash Buffer (0.2M Tris-HCl pH9, 0.2M KCl, 0.025M EGTA pH8, 0.035M MgCl_2_, 1 mM DTT, 1 mM PMSF, 100 ug/ml cycloheximide, 50 ug/ml chloramphenicol). Wash was repeated twice before resuspending the beads in 1 mL of wash buffer, transferring the sample to a 1.5 mL tube, and repeating the wash three times using 1 mL of buffer. Washed beads were resuspended in 800 uL of lysis binding buffer (100 mM Tris-HCl pH8, 1M LiCl, 10 mM EDTA pH8, 1% (v/v) SDS, 5 mM DTT, 1.5% (v/v) Antifoam A) and frozen at -80°C until mRNA isolation.

mRNAs were isolated from both the Total and TRAP fractions using biotinylated oligo(dT) and streptavidin magnetic beads as previously described [100]. Purified mRNAs were processed into libraries as described above, omitting the DNase I treatment.

### RNA- and TRAP-Sequencing Data Processing

Files were trimmed using cutadapt [102] and quality reports were generated using FastQC [103]. Following processing steps were completed using the systemPipeR package [104]. Trimmed reads were mapped to the TAIR10 Arabidopsis genome using HISAT2 v2.2.1 [105]. The number of sequenced and aligned reads is provided in Table S1. The summarizeOverlaps function from the GenomicRanges package [106] was used to get read counts using the Araport11 gff (201606) [107]. Normalized rlog values were generated with the DESeq2 package. Expression data are provided in Dataset S1 and for the total and TRAP-sequencing in Dataset S6.

PCA plots were generated on rlog normalized data using the prcomp function in R. Four samples (*cca1lhy* 37°C ZT15 RepA, *cca1lhy* 37°C ZT3 RepB, WT 22°C ZT15 RepB, and WT 22°C ZT18 RepB) were excluded from the transcriptome analyses as they largely varied from the other two biological replicates for that condition and thus were excluded from all other downstream analyses.

#### Cycling and Rhythmicity Analysis

Circadian oscillations/cycling genes were determined separately for WT and *cca1lhy* in the time-course transcriptome data using Metacycle and the meta2d function [108]. Genes with <20 reads were excluded from the analysis. A cutoff of BH.Q <0.05 was used to determine significantly cycling transcripts (Dataset S1). Differences in cycling were determined using the R package LimoRhyde filtering for adj. P<0.05 [109]. Genes cycling in our WT data were compared to cycling genes from other time-course RNA-sequencing or microarray datasets performed in seedlings of approximately the same age where they were entrained in constant temperature (22°C) and 12 light:12 dark photocycles before being transitioned to constant light (LL) for two days before collection [21,34,53,54]. Cycling genes called by each dataset as well as the overlapping core cycling genes are provided in Dataset S1. Similarly, genes that did not meet the cycling threshold in any of the datasets used were compiled and provided in Dataset S1.

#### Phase, timing of peak gene expression and amplitude estimation

Difference in time of peak expression was determined, for all genes that cycle in WT by finding their time of peak expression based on read counts (not using phase values from the Metacycle analysis). As *cca1lhy* mutant lines have been previously shown to have an advanced circadian phase, the difference in time of peak expression was calculated as Time*_cca1lhy_-*Time_WT_ [12,51]. We considered positive values as a phase delay and converted them to the equivalent 24 h circular shift of phase advance. Cycling amplitudes were calculated by finding 1/2 the difference in rlog values between the highest and lowest expression for each genotype ((max-min)/2).

#### Likelihood Ratio Test

To identify genes with altered expression in *cca1lhy*, a likelihood ratio test (LRT) was performed to determine the effects of genotype and time using the DESeq2 package, and on genes with >20 read counts [110]. The full model was: design = ∼Time + Temperature + Genotype + Time:Temperature + Time:Genotype + Temperature:Genotype + Time:Temperature:Genotype, and significant interactions were considered as FDR<0.05 (Dataset S3). LRTs using the structure ∼Time+Temperature+Time:Temperature were used to identify gated genes within each genotype, with genes with a Time:Temperature effect considered gated. Significant results of both the pooled and split genotype LRTs are provided in Dataset S3. A LRT with the model ∼Time+Genotype+Time:Genotype was also run on only ambient samples, with the results in Dataset S2.

#### Motif enrichment analysis

Motifs were identified in gene promoters (defined as 1 kb upstream to 100 bp downstream of the TSS based on Araport11 gene annotations) using the vmatchPattern function from the Biostrings package with exact matching [111]. Evening Elements (EEs) were defined as AAATATCT and CCA1-binding sites (CBS) as AAAAATCT.

#### Differential Expression Analysis

Differentially expressed genes (DEGs) were determined using DESeq2 by comparing between heat and control conditions within each genotype at each timepoint. Genes with FDR <0.05 and LFC >1 were considered as a DEG (Dataset S3). DESeq2 was also used to compare between genotypes at each timepoint in each treatment for comparison to the proteome. For this comparison, genes with an FDR <0.05 were considered significantly different between genotypes (Dataset S2 and S3).

#### Magnitude of Heat Response

Magnitude of heat responses was calculated for genes with altered temperature responses in *cca1lhy.* LFC values comparing heat (37°C) and control (22°C) samples within each genotype were extracted from DESeq2 analysis and averaged across timepoints. Genes were considered as either up- or -down regulated based on the sign (positive or negative) of the LFC. Unpaired Welch’s t-tests were used to identify differences between genotypes using P <0.05.

### GO Enrichment and Co-expression Analysis

Gene Ontology (GO) enrichment was determined using Panther Biological Function, using a Fisher’s exact test and calculating the FDR. Terms with FDR <0.05 were considered significant [112]. For generating graphs, term lists were simplified as previously described using Revigo [113,114]. Full GO term lists are provided in Dataset S4.

Co-expression analyses were generated as described in Wisecaver et al. 2017 using scripts provided in their GitHub [115,116]. Briefly, Pearson’s correlation coefficients were calculated for each gene pair in the data, which is used to determine a mutual rank score. Mutual rank scores were then converted into edge weights with 3 different decay functions, each producing a distinct network with varying stringency for connections. Finally, co-expressed gene sets were generated using ClusterONE [117]. Four clusters from decay co-efficient 10 containing *HSFA1E* were identified and all members of the clusters were plotted using Cytoscape [118].

### Data Visualization

Most plots were generated in R using the ggplot2 R package. Upset plots were generated using the UpSetR package [119]. Heatmaps were generated using the R package “pheatmap”. Some figures were generated using GraphPad Prism 10. Figures were assembled in Affinity.

#### Dual Luciferase Assay in *Nicotiana*

The *UBA2A* promoter (313 bp), defined as from the ATG to the nearest upstream gene, was cloned into pGreenII 0800-LUC by digesting with HindIII and NcoI followed by ligation with T4 DNA Ligase (Table S3). The coding sequences of CCA1 and LHY were cloned into pGWB505 destination vectors to generate 35S:CCA1-GFP and 35S:LHY-GFP. A 35S:TurboID-YFP plasmid was used as a control. The promoter and effector constructs were co-infiltrated into ∼ 4 - 5 week old *Nicotiana benthamiana* plants (tobacco). After 2 d, leaf discs from two independently infiltrated leaves were pooled to produce 1 sample. Tissue was then snap-frozen and ground. The dual luciferase assay was then performed using the Promega Dual Luciferase Reporter Assay System. Briefly, tissue was re-suspended in 100 ul of 1X Passive Lysis Buffer, incubated for 15 min on ice and then centrifuged for 15 min at 4°C 18,000 g before transferring the supernatant. Equal volumes (100 ul) of LarII and Stop & Glo were then used. Luminescence was read using a Promega GloMax Multiplus Plate Reader (dual injectors).

#### Liquid Chromatography-Tandem Mass Spectrometry (LC-MS/MS) protein extraction and Trypsin digestion

WT and *cca1lhy* seedlings were grown and collected as described above. Ground, lyophilized samples were resuspended in extraction buffer (50 mM Tris-HCl pH 8.0, 4% (w/v) SDS) at a sample:buffer ratio of 1:2 (w/v). Samples were boiled at 95°C with 1500 rpm shaking (Eppendorf ThermoMixer F2.0) for 10 min. Subsequently, centrifugation was performed at 20,000 x g for 10 min to clarify samples and supernatant was moved to a fresh Eppendorf tube. The supernatant was reduced with 10 mM DTT for 10 min at 95°C followed by alkylation with iodoacetamide for 30 min at room temperature in the dark without shaking. Peptide pools were generated using automated sample processing on a KingFisher APEX (Thermo Fisher Scientific) and proteins were digested overnight with sequencing-grade trypsin (Cat# V5113; Promega) diluted in 50 mM triethylammonium bicarbonate buffer (TEAB) pH 8.5 (T7408; Sigma). Following digestion, samples were acidified using trifluoroacetic acid (TFA; Cat# A117; Fisher) to 0.5% (v/v) final concentration. Peptides were desalted using an OT-2 liquid handling robot (Opentrons Labworks Inc) with Omix C18 pipette tips (A5700310K; Agilent). The final desalted peptides were dried in a speedvac (Thermo Fisher Scientific) at room temperature and stored at -80℃ until use. Prior to mass spectrometry analysis, peptides were resuspended in 3% (v/v) acetonitrile / 0.1% (v/v) formic acid.

#### Protein LC-MS/MS Processing

One µg of resuspended peptide was eluted into the mass spectrometer using an HT µPac Neo Plus column (Thermo Fisher Scientific; COL-UPAC005CAP) using -30, -50, and -70 CVs. Peptides were analyzed using a FAIMSpro enabled Fusion LUMOS Orbitrap mass spectrometer (Thermo Fisher Scientific) in BoxCarDIA mode as previously described [120,121]. A spray voltage of 2.2 kV, funnel RF level of 40 and heated capillary at 300°C was employed, with all data acquired in profile mode using positive polarity with peptide match off and isotope exclusion selected. Gradients were run at .12 µL/min and analytical column temperature at 50°C. A solvent B gradient (0.1% (v/v) FA in 80% (v/v) ACN) was deployed stepwise from 3 - 18% (0 –17 min), 18 - 28% (17 – 24 min), 28 - 45% (24 – 27 min) and 45% - 100% (27 – 30 min) to elute peptides. MS1 analysis was performed using two multiplexed targeted SIM scans of 10 BoxCar windows each, with a resolution of 120,000 for detection at 200 m/z, with normalized AGC targets of 100% per BoxCar isolation window. Windows were custom designed as previously described [120]. An AGC target for MS2 fragment spectra was set at 2000%. Twenty-eight 38.5 m/z windows were used with 1 m/z of overlap. The resolution used was 30,000 using dynamic maximum injection time and 6 was used as the minimum number of desired points across each peak.

#### Protein LC-MS/MS Data Analysis

BoxCarDIA data was analyzed in Spectronaut v20.6 (Biognosys AG) using a library-free DIA approach under default settings. Data were searched using the Araport 11 proteome (Phytozome) [122]. Key search parameters were as follows: trypsin digest allowing two missed cleavages, fixed modifications (carbamidomethyl [C]), variable modifications (oxidation [M]), a peptide spectrum match (PSM), and peptide and protein false discovery threshold of 1%. Data was Log2 transformed, median normalized and significantly changing differentially abundant proteins (DAPs) were determined and corrected for multiple comparisons (Bonferroni-corrected *p* value <0.05; *q* value, LFC >1). Magnitude of heat responses was assessed by averaging LFC across the two timepoints within each genotype. GO enrichment was performed as described above, using only the first protein annotated if the protein group reflects multiple proteins.

Patterns of behavior between the transcriptome and proteome were assessed using comparisons between the genotypes after heat stress at both ZT48 and ZT60 (*cca1lhy* 37°C vs WT 37°C) (Datasets S2, S3, and S5). Analyses were performed using DESeq2 or Spectronaut as specified above applying a FDR or *q* value <0.05 and no LFC threshold. For ambient comparisons, the comparison was subset to genes with a genotype effect based on the LRT on ambient expression in the RNA-seq. For heat-treated conditions, the comparison was subset to genes with a genotype effect based on the LRT on the temperature response in the RNA-seq.

#### Metabolomics Sample Preparation

WT and *cca1lhy* seedlings were grown, and four biological replicates were collected as described above. Samples were snap-frozen and then freeze-dried for 3 days. Tissue was then ground in a chilled bead-beater (Omni Bead Ruptor 24). Approximately 10 mg of powder was weighed in glass vials and extracted with 100 ul/mg of monophasic extraction solvent (30:30:20:20 acetonitrile, methanol, isopropanol and water). Samples were extracted by sonication on ice for 5 min followed by 90 min vortexing at 4°C. The samples were centrifuged at 4°C for 30 min at 3000 g. Supernatants were collected and pipetted into LC-MS vials. A quality control sample was constituted by combining equal volumes of all samples and used to monitor system stability.

#### Metabolomic LC-MS methods

Targeted metabolomics of polar, primary metabolites was performed at the UC Riverside Metabolomics Core Facility as described previously [123,124]. Briefly, analysis was performed on a TQ-XS triple quadrupole mass spectrometer (Waters) coupled to an I-class UPLC system (Waters). Separations were carried out on a ZIC-pHILIC column (2.1 x 150 mm, 5 µM) (EMD Millipore). The mobile phases were (A) water with 15 mM ammonium bicarbonate adjusted to pH 9.6 with ammonium hydroxide and (B) acetonitrile. The flow rate was 200 µL/min and the column was held at 50°C. The injection volume was 2 µL. The gradient was as follows: 0 min, 90% B; 1.5 min, 90% B; 16 min, 20% B; 18 min, 20% B; 20 min, 90% B; 28 min, 90% B.

The MS was operated in multiple reaction monitoring mode (MRM). Source and desolvation temperatures were 150°C and 600°C, respectively. Desolvation gas was set to 1100 L/hr and cone gas to 150 L/hr. Collision gas was set to 0.15 mL/min. All gases were nitrogen except the collision gas, which was argon. Capillary voltage was 1 kV in positive ion mode and 2 kV in negative ion mode. A quality control (QC) sample, generated by pooling equal aliquots of each sample, was analyzed periodically to monitor system stability and performance. Samples were analyzed in random order.

#### Metabolomic LC-MS Data processing and Analysis

Targeted data processing was performed with the open-source Skyline software [125]. After metabolite identification and relative quantification, we used a UCR Metabolomics Core proprietary tool, HTML based solution for data handling. Principal Components Analysis (PCA) was performed using the R function prcomp on log2 transformed data with scale=TRUE [126]. The heatmap was generated using z-scores calculated on log2 transformed relative abundance values. For pairwise comparison between treatments, Welch’s t-test was applied to Log2 transformed data. P-values were corrected by using the Benjamini and Hochberg (1995) correction (Dataset 7) [127]. For specific examples, metabolites significantly differing between *cca1lhy* and WT and temperature treatments were analyzed using a 2-way ANOVA on Log2 transformed data.

#### Statistical analysis

All statistical analyses were performed using the program R v4.4.2.

## Acknowledgements

The authors thank Samantha Cordingley and Gabriela Brown for critical reading of the manuscript. We also thank Dr. Julia Bailey-Serres’s lab for constructive feedback and discussions. We thank Dr. Dmitri Nusinow for providing pGreenII 0800-LUC plasmid.

## Authors Contribution

D.H.N and R.I.S conceived the experiments. R.I.S and D.H.N. performed the experiments and analysis, excluding the proteomics, which was performed by L.E.G and R.G.U. R.I.S and D.H.N wrote the manuscript. All authors approve the manuscript.

## Data Availability

All transcriptomic data reported in this manuscript are accessible from the Gene Expression Omnibus (GEO) database GSE354530. All raw proteomics data can be found through the Proteomics IDEntifications Database (PRIDE; https://www.ebi.ac.uk/pride/) using the dataset identifier PXD087269. Gene Accession numbers mentioned in this study can be found in Table S2.

**Supplementary Figures and Tables:** Uploaded as a separate document.

**Supplementary Datasets**: Available upon request.

