## Supplementary Figures and Tables for "Circadian clock components CCA1 and LHY shape heat responses across regulatory layers"

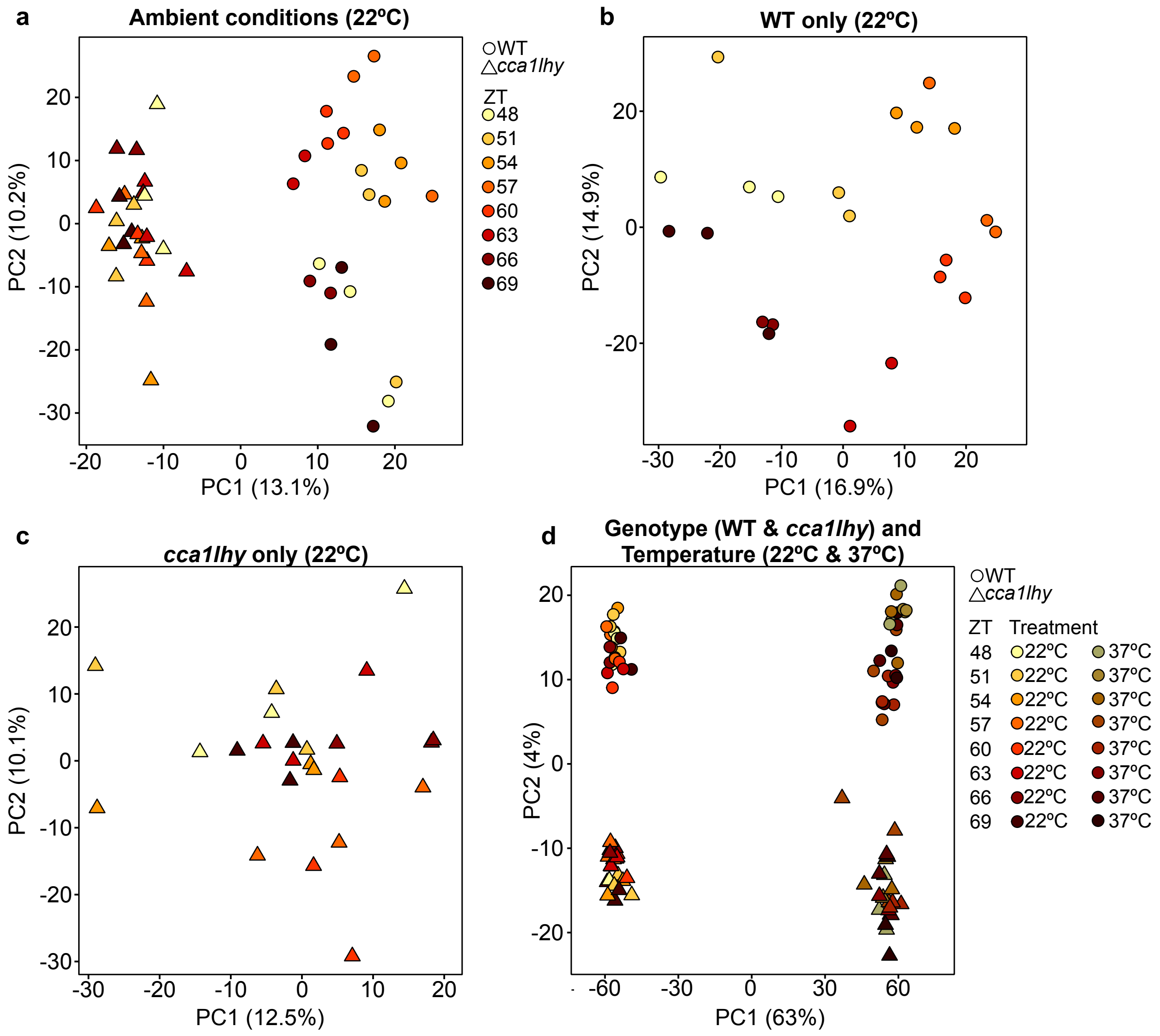


**Figure S1:** Data separation in the RNA time-course is explained by genotype, temperature, and time. **a)** Principal component analysis (PCA) of all quantified genes in ambient conditions (22°C). **b)** PCA of all quantified genes in WT at 22°C **c)** PCA of all quantified genes in *cca1lhy* at 22ºC **d)** PCA of all quantified genes in *cca1lhy* and WT from the 22°C and 37°C (1 h) treated samples from the RNA time-course. Color indicates time and treatment in **d**. Shape indicates genotype with circles representing WT and triangles representing *cca1lhy*.

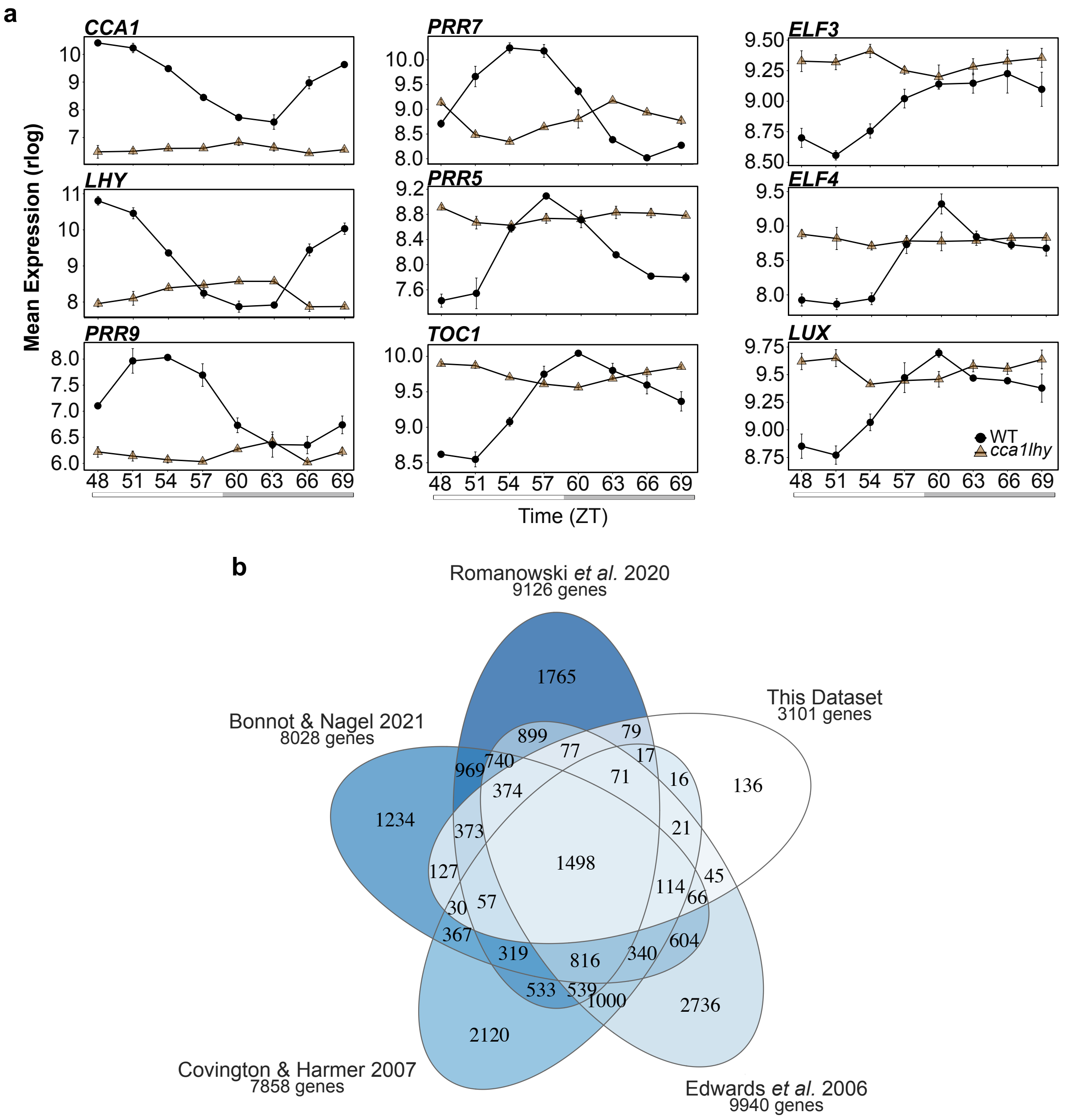


**Figure S2:** Cycling genes in WT are mis-regulated in *cca1lhy*. **a)** Core clock gene expression at 22ºC. Transcript profiles are displayed as average rlog +/- SE. White bars represent subjective day; grey bars represent subjective night. **b)** Overlap of cycling genes from RNA-seq or microarray circadian time-course datasets performed on seedlings entrained in light/dark (LD) cycles (12 h light: 12 h dark) and constant 22°C that had been transitioned to constant light (LL) for 2 days [21,34,53,54].

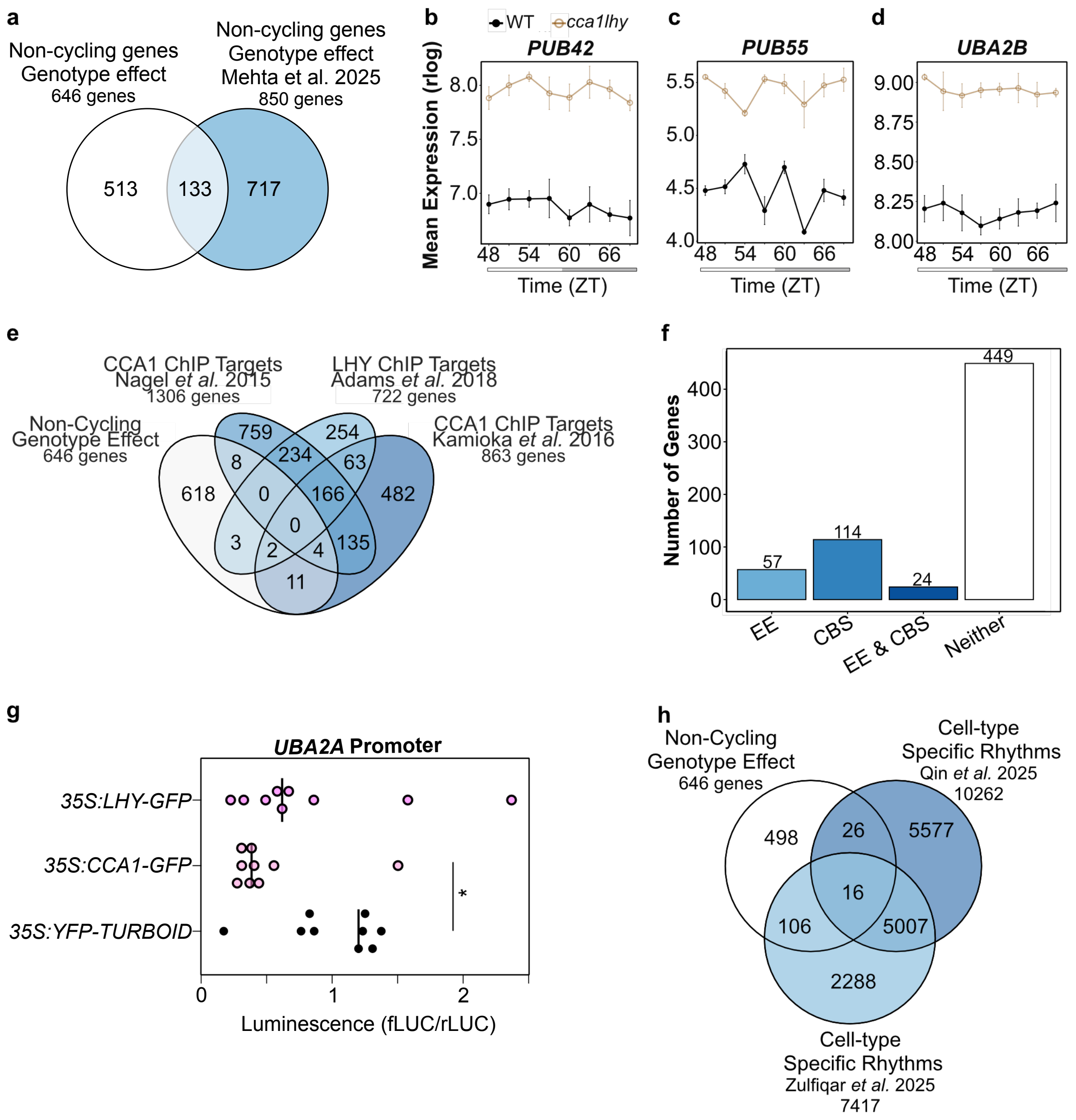


**Figure S3:** A subset of non-cycling genes regulated by *CCA1* and *LHY* cycle in single-cell datasets. **a)** Overlap of non-cycling genes with a genotype effect between our dataset and Mehta et al. 2025 [52] **b)** Ambient expression profile of *PUB42,* **c)** *PUB55,* and **d)** *UBA2B.* **e)** Non-cycling genes with altered expression in *cca1lhy* and their overlap with published ChIP-sequencing targets of CCA1 and LHY [11–13]. **f)** Number of non-cycling genes with altered expression in *cca1lhy* that contain an EE (Evening Element; AAATATCT), CBS (CCA1-binding site;

AAAAATCT), both, or neither motif. **g)** Dual luciferase reporter assay of *UBA2A:LUC* activity. Data normalized so that control expression is set to 1. Each dot represents a biological replicate. Student's t-test *p<0.05. **h)** Overlap between non-cycling genes mis-regulated in *cca1lhy* and time-course single-cell experiments [59,60].

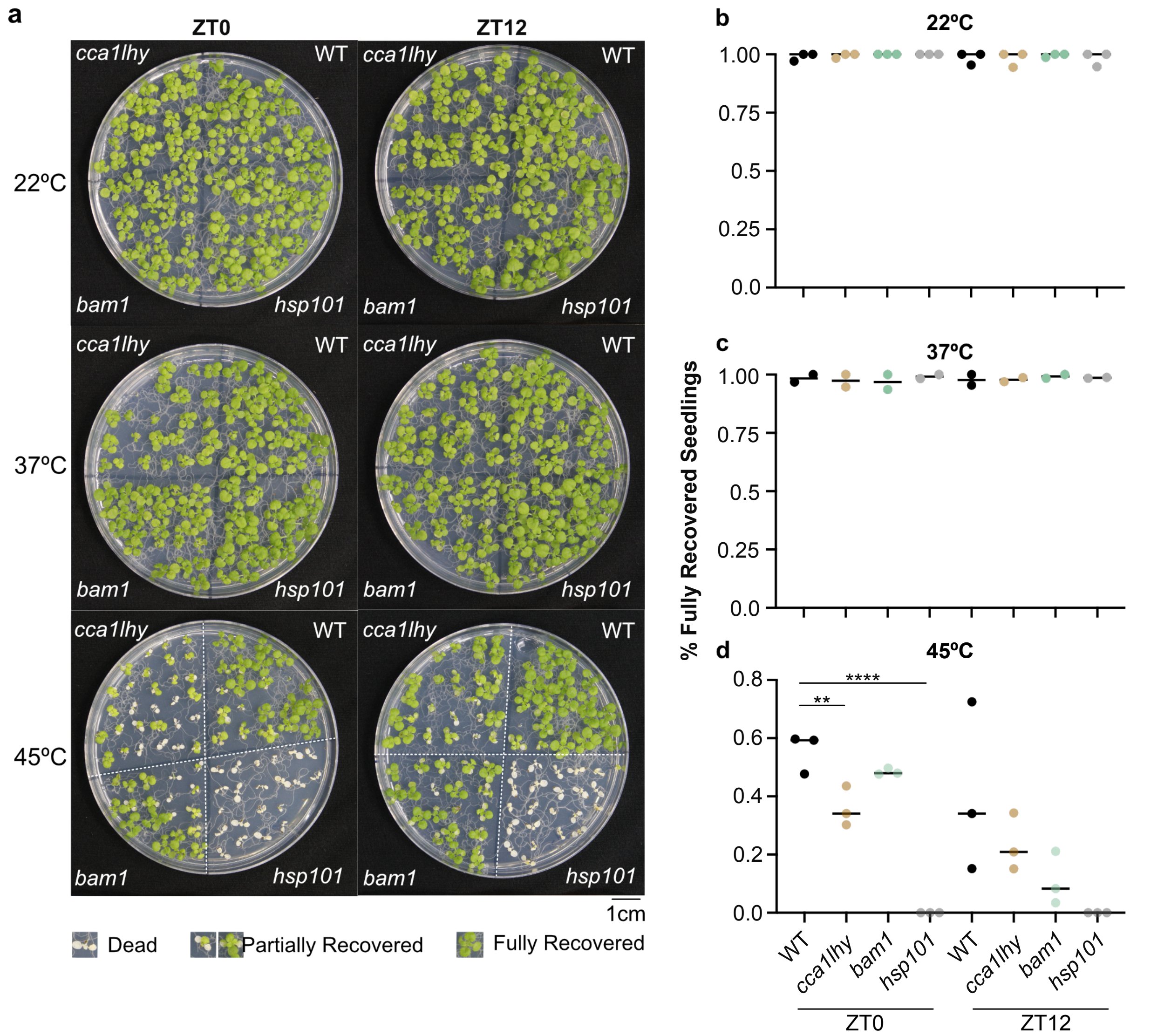


**Figure S4:** *cca1lhy* has reduced survival after heat stress. **a)** Representative images. 22°C indicates plates that remained in control conditions (22°C) for the duration of the experiment. 37°C plates experienced 30 min 37°C before being returned to control (22°C). 45°C plates experienced 30 min 37°C, 30 min 22°C, 90 min 45°C before being returned to control (22°C). Images were taken after a 6-day recovery period. **b)** Quantification of fully recovered seedlings on 22°C plates (N=3), **c)** 37°C plates (N=2), and **d)** 45°C plates (N=3). $^{**}p<0.01$, $^{****}p<0.0001$.

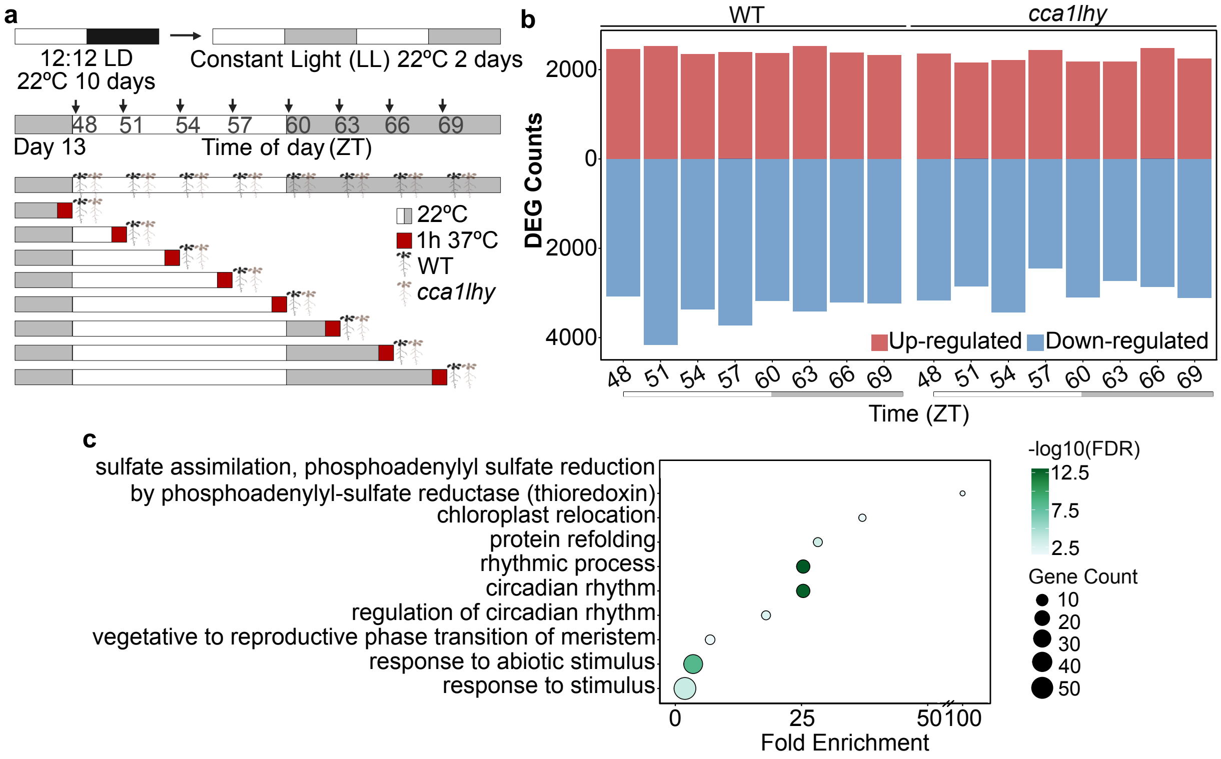


**Figure S5:** Genes mis-regulated in *cca1lhy* after a heat treatment. **a)** Experimental design. Grey boxes represent the subjective night. Red indicates a 1 h 37°C treatment. **b)** DEGs called at each timepoint in each genotype divided by up and down-regulated genes. White and grey boxes represent subjective day and night respectively. **c)** Enriched GO processes for cycling genes with a genotype effect on their temperature responses.

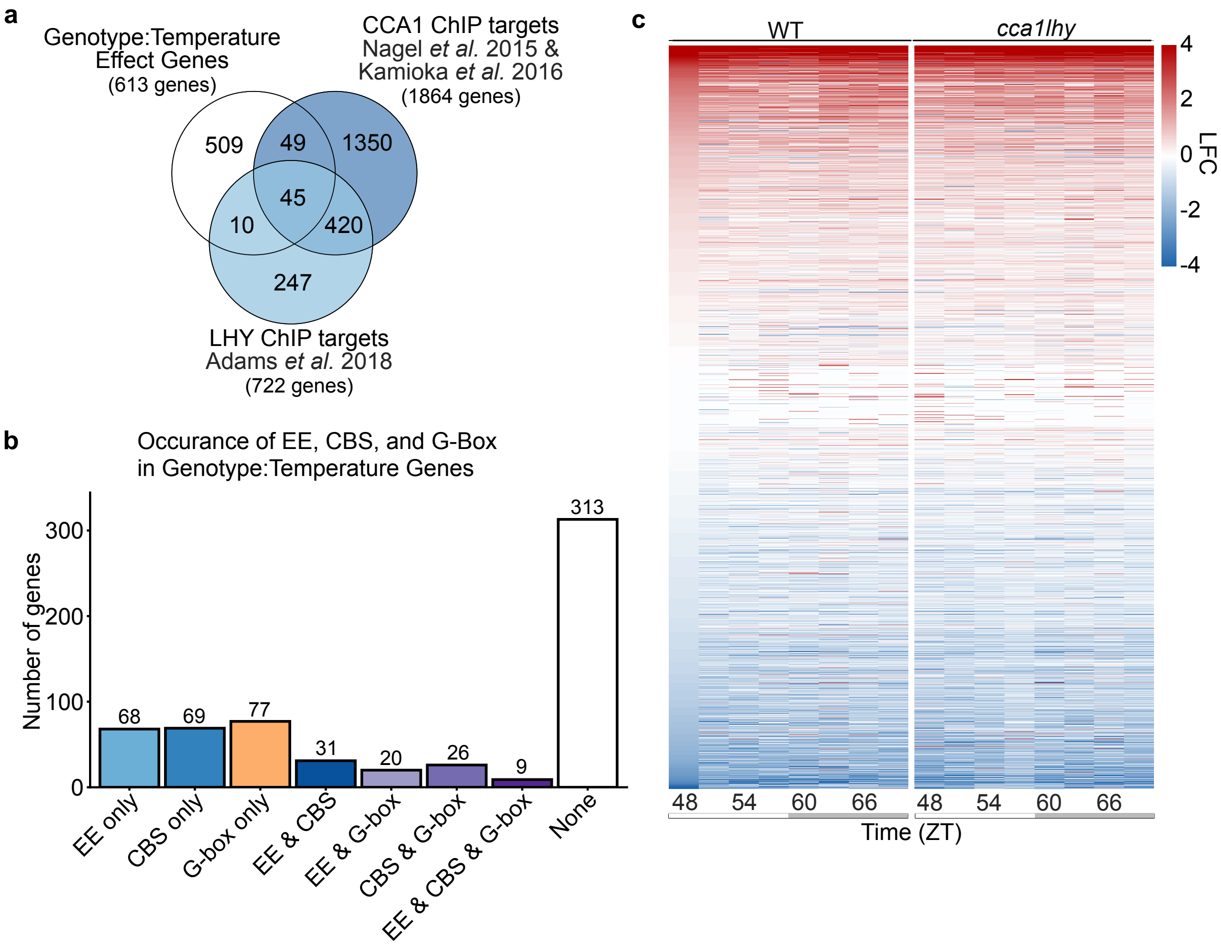


**Figure S6:** CCA1 and LHY may directly regulate a subset of genes with a genotype:temperature effect. **a)** Overlap between genes with altered heat responses in *cca1lhy* and CCA1 and LHY ChIP-sequencing targets. **b)** Occurrence of Evening Element (EEs) or CCA1 Binding Site (CBS) motifs in genotype:temperature effect genes. **c)** Heatmap of the LFC of all CCA1 or LHY ChIP-sequencing targets. Genes ordered by LFC at ZT48 in WT. Grey bars under the x-axis indicate subjective night.

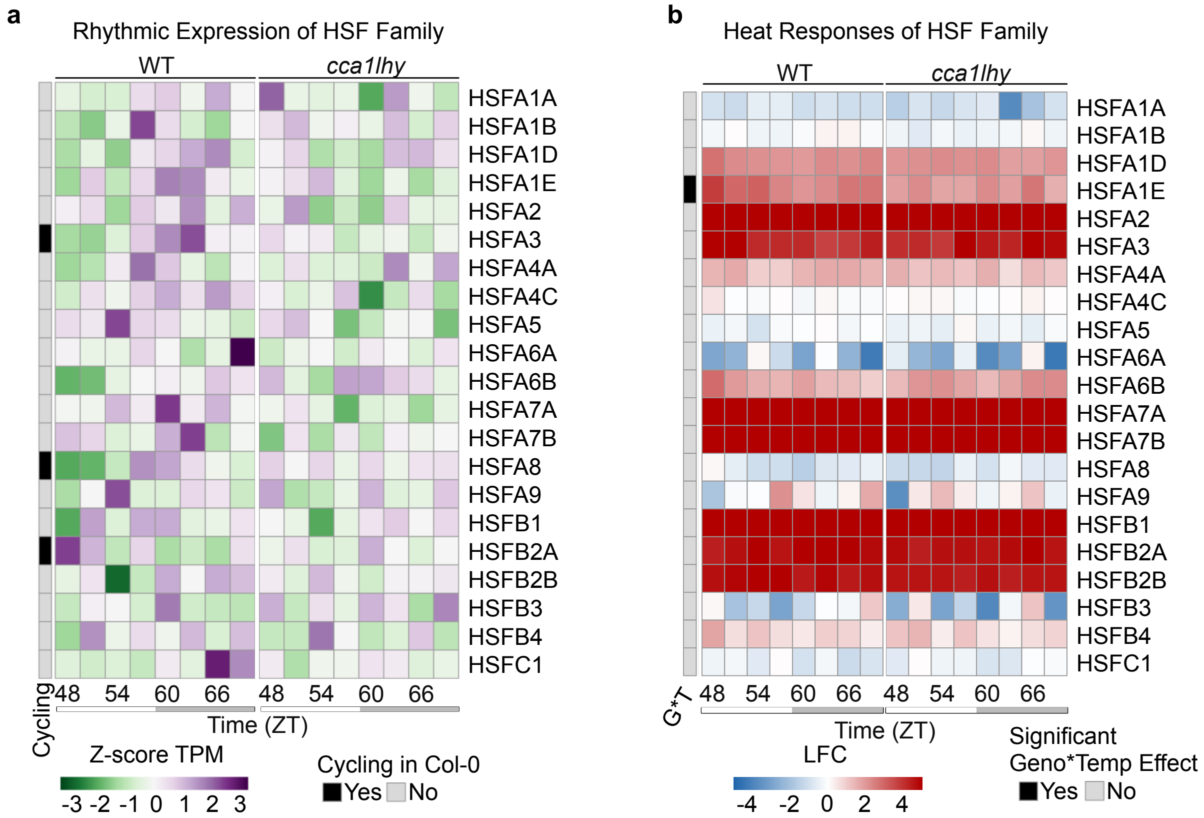


**Figure S7:** Heat Shock Transcription Factors (HSFs) have altered expression in *cca1lhy*. **a)** Transcript profiles of all HSFs in Arabidopsis in ambient conditions (22°C). Data shown as normalized TPM values. Genes that cycle in WT indicated by black boxes. **b)** Heat responses of all HSFs. Data shown as LFC between heat (37°C) and control (22°C) samples. Genes with a significant genotype effect on temperature response based on the Likelihood Ratio Test (LRT) indicated by black boxes. White and grey boxes represent subjective day and night respectively.

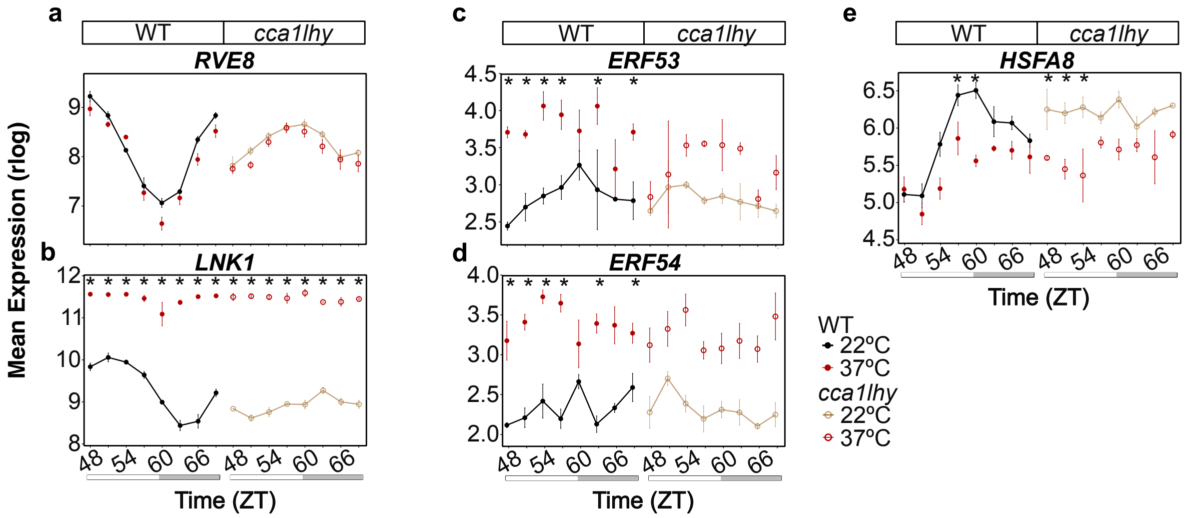


**Figure S8:** Mis-regulation of clock genes linked to thermotolerance and their targets. **a)** Transcript profiles of *RVE8*, **b)** *LNK1*, **c)** *ERF53*, **d)** *ERF54*, and **e)** *HSFA8*. White and grey boxes under the axis represent subjective day and night respectively. Asterix (*) represents DEG (FDR <0.05, LFC >1) at the indicated timepoint.

**Figure S9:** Altered protein abundance in *cca1lhy* in ambient conditions (22°C). **a)** Differentially abundant proteins (DAPs) between *cca1lhy* and WT at each timepoint and their overlap. **b)** DAPs within *cca1lhy* or WT between the two timepoints (ZT48 and ZT60). **c)** Comparison of LFC (*cca1lhy* vs WT) in the transcriptome and proteome in ambient conditions (22°C) at ZT48 and at **d)** ZT60. Genes with altered expression in *cca1lhy* in the transcriptome are plotted. Purple dots (circles) are genes significantly upregulated in *cca1lhy,* and green dots (circles) are significantly downregulated in *cca1lhy* in both the transcriptome and proteome. Black dots (circles) are upregulated in either the transcriptome or proteome and downregulated in the other. **e)** Expression of concordant genes in Mehta *et al.* 2026 [52]. Only concordant genes at ZT48 in this dataset were compared to the Mehta *et al.* data at ZT1 and the same for ZT60/ZT13. Purple indicates gene is upregulated in both the transcriptome and proteome in this dataset, green indicates gene is downregulated in both the transcriptome and proteome. **f)** Transcript and protein profiles of UBA2A and **g)** CSU4. Asterisk (*) indicates significant DEG or DAP based on FDR (transcriptome) or q (proteome) $<0.05$.

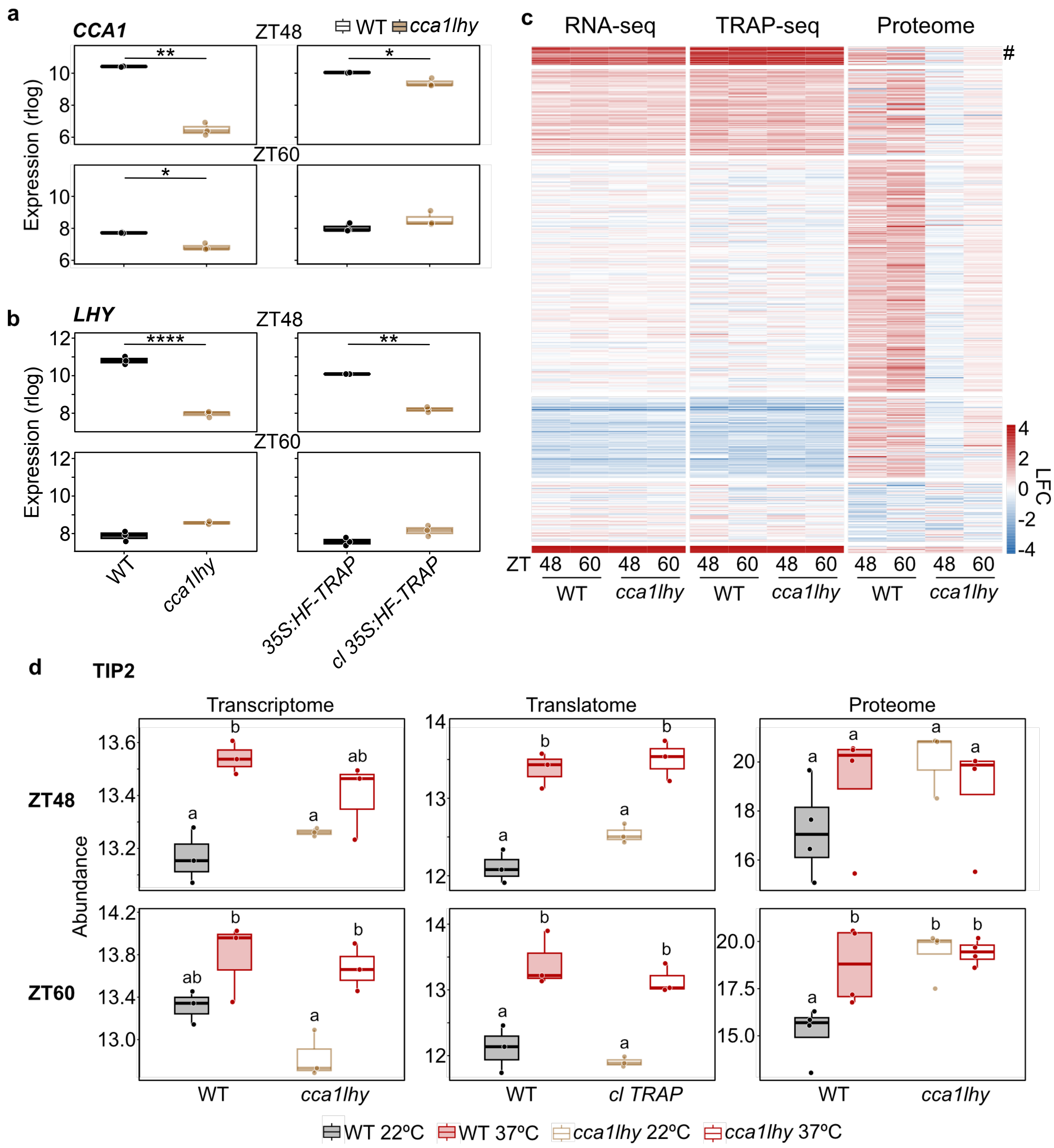


**Figure S10:** Heat-responsive post-translational regulation may be altered in *cca1lhy*. **a)** and **b)** Expression of *CCA1* (a) or *LHY* (b) in WT, *cca1lhy, 35S:HF-RPL18*, and *cca1lhy (cl) 35S:HF-RPL18*. Unpaired t-test *p<0.05, **p<0.01, ****p<0.0001. **c)** K-means clustering of the LFC of DAPs specific to one genotype across the transcriptome, translatome, and proteome. Number (#) indicates the location of the TONOPLAST INTRINSIC PROTEIN 2 (TIP2) and PLASMA MEMBRANE INTRINSIC PROTEINs (PIPs) discussed in the main text. **d)** Example of TIP2, which displays genotype-specific post-translational regulation. Abundance is represented by rlog in the RNA and TRAP-sequencing and by log2 relative abundance in the proteome. Significant differences based on a 2-way ANOVA with Tukey's significant difference test indicated by lowercase letters ($p<0.05$).

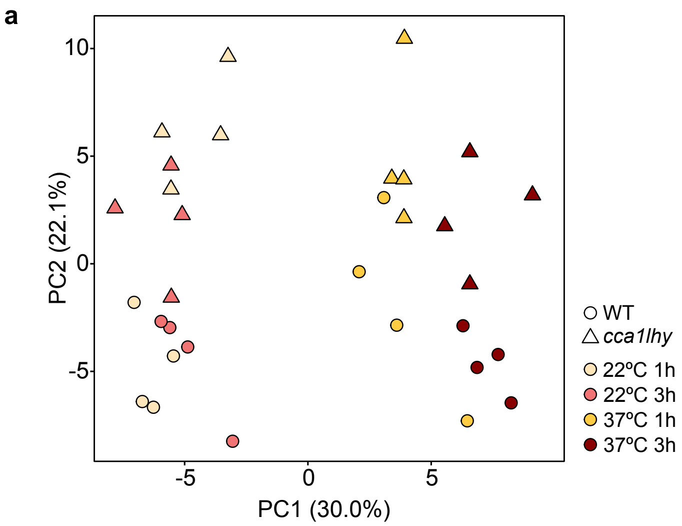


**Figure S11:** Metabolomic samples cluster by temperature and genotype. **a)** PCA of metabolomic samples. 1 h represents samples treated from ZT47-48, 3 h samples were treated from ZT47-ZT50.

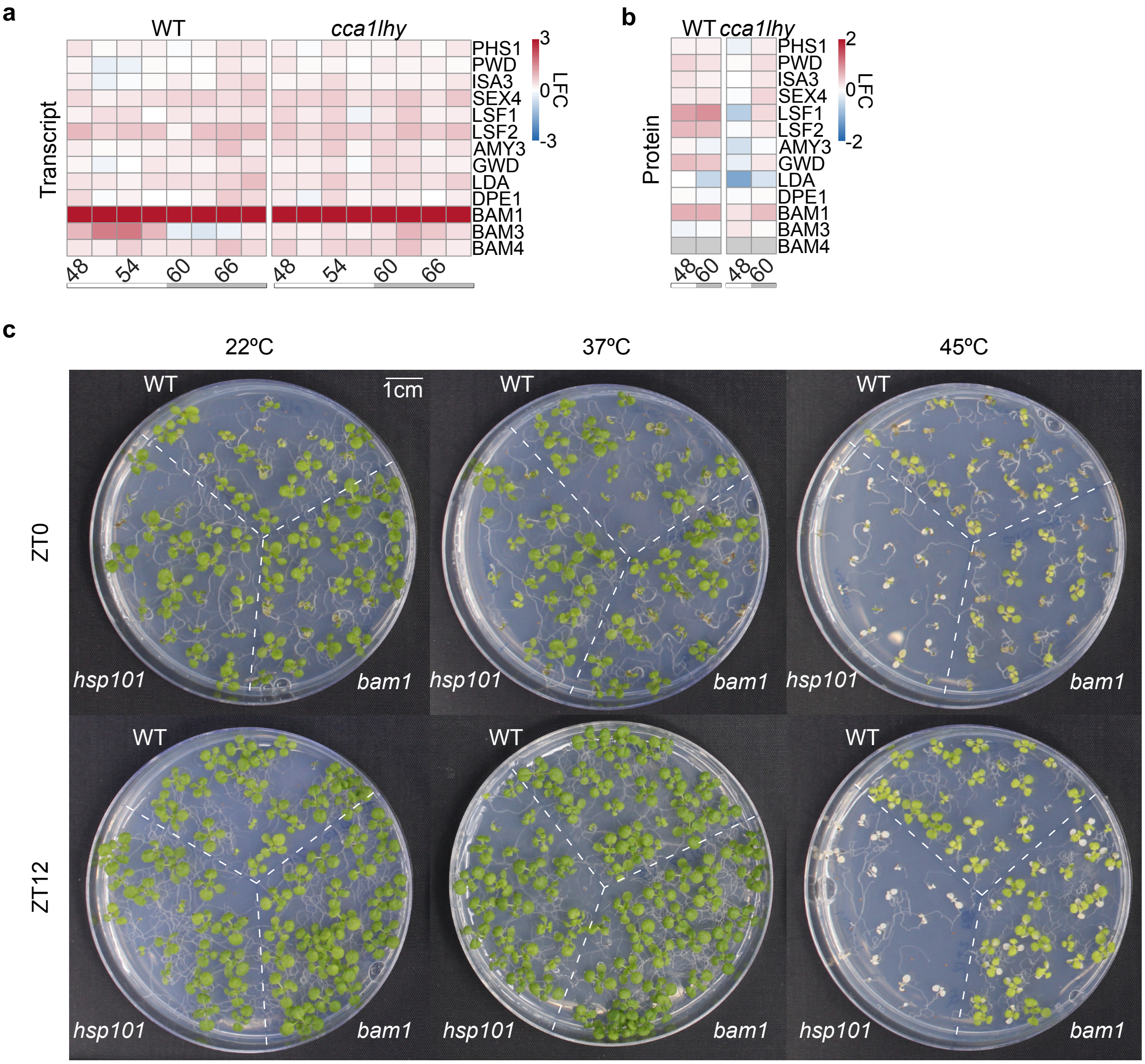


**Figure S12:** *BAM1* mutants may have altered thermotolerance. **a)** LFC of genes involved in starch degradation in response to heat (1 h 37°C) in the transcriptome and **b)** the proteome. Grey bars represent the subjective night. **c)** Representative images of acquired thermotolerance assays, performed as shown in Fig. 2. 22°C and 37°C plates are controls. Scale bar is 1cm. Quantification provided in Fig. 5j.

**Table S1:** Read mapping percentages for RNA-seq libraries

| SAMPLE | Nreads2x | Nalign | Perc_Aligned | Nalign_Primary | Perc_Aligned_Primary |
|---|---|---|---|---|---|
| A0CC | 57770152 | 44879903 | 77.6870087 | 44879903 | 77.6870087 |
| A0CH | 11440038 | 10423570 | 91.1148197 | 10423570 | 91.1148197 |
| A0WC | 70983472 | 58118524 | 81.8761359 | 58118524 | 81.8761359 |
| A0WH | 13322516 | 10997515 | 82.548334 | 10997515 | 82.548334 |
| A3CC | 10252178 | 8128020 | 79.2809099 | 8128020 | 79.2809099 |
| A3CH | 32807650 | 25100652 | 76.5085338 | 25100652 | 76.5085338 |
| A3WC | 12163998 | 11119795 | 91.4156267 | 11119795 | 91.4156267 |
| A3WH | 45163386 | 37540182 | 83.1208316 | 37540182 | 83.1208316 |
| A6CC | 43473158 | 37759721 | 86.8575524 | 37759721 | 86.8575524 |
| A6CH | 23163502 | 18878987 | 81.5031639 | 18878987 | 81.5031639 |
| A6WC | 15572776 | 13402814 | 86.06567 | 13402814 | 86.06567 |
| A6WH | 13121044 | 10655028 | 81.2056419 | 10655028 | 81.2056419 |
| A9CC | 41009988 | 34918644 | 85.1466818 | 34918644 | 85.1466818 |
| A9CH | 30142922 | 24283907 | 80.5625513 | 24283907 | 80.5625513 |
| A9WC | 12656246 | 11036756 | 87.2040256 | 11036756 | 87.2040256 |
| A9WH | 39308766 | 33717950 | 85.7771775 | 33717950 | 85.7771775 |
| A12CC | 16476166 | 14474824 | 87.8531085 | 14474824 | 87.8531085 |
| A12CH | 25264566 | 19958007 | 78.9960413 | 19958007 | 78.9960413 |
| A12WC | 14389574 | 12327620 | 85.6705 | 12327620 | 85.6705 |
| A12WH | 17610054 | 14346742 | 81.4690404 | 14346742 | 81.4690404 |
| A15CC | 20161568 | 18566973 | 92.0909177 | 18566973 | 92.0909177 |
| A15CH | 23412584 | 17762633 | 75.867888 | 17762633 | 75.867888 |
| A15WC | 16093894 | 12727472 | 79.0826136 | 12727472 | 79.0826136 |
| A15WH | 28047642 | 22018706 | 78.50466 | 22018706 | 78.50466 |
| A18CC | 19242312 | 16748151 | 87.0381428 | 16748151 | 87.0381428 |
| A18CH | 14963046 | 12967193 | 86.6614525 | 12967193 | 86.6614525 |
| A18WC | 62452626 | 51474222 | 82.4212292 | 51474222 | 82.4212292 |
| A18WH | 39417114 | 32363921 | 82.1062673 | 32363921 | 82.1062673 |
| A21CC | 48447578 | 36778038 | 75.913058 | 36778038 | 75.913058 |
| A21CH | 22316824 | 17216043 | 77.1437862 | 17216043 | 77.1437862 |
| A21WC | 18770126 | 15568195 | 82.9413452 | 15568195 | 82.9413452 |
| A21WH | 10996626 | 9032554 | 82.1393216 | 9032554 | 82.1393216 |
| B0CC | 7796192 | 6687105 | 85.7739907 | 6687105 | 85.7739907 |
| B0CH | 11461984 | 9586541 | 83.6377105 | 9586541 | 83.6377105 |
| B0WC | 9083736 | 8015194 | 88.2367563 | 8015194 | 88.2367563 |
| B0WH | 14825676 | 11768626 | 79.3800296 | 11768626 | 79.3800296 |

| | | | | | |
|---|---|---|---|---|---|
| B3CC | 10967788 | 8842732 | 80.6245708 | 8842732 | 80.6245708 |
| B3CH | 14814740 | 10378913 | 70.0580165 | 10378913 | 70.0580165 |
| B3WC | 10494730 | 8282605 | 78.9215635 | 8282605 | 78.9215635 |
| B3WH | 128299638 | 94582364 | 73.7198994 | 94582364 | 73.7198994 |
| B6CC | 7354808 | 6240842 | 84.8539078 | 6240842 | 84.8539078 |
| B6CH | 18588882 | 15345181 | 82.5503169 | 15345181 | 82.5503169 |
| B6WC | 11119520 | 8401656 | 75.5577219 | 8401656 | 75.5577219 |
| B6WH | 17185542 | 13903028 | 80.8995608 | 13903028 | 80.8995608 |
| B9CC | 22185156 | 17630359 | 79.4691685 | 17630359 | 79.4691685 |
| B9CH | 16529908 | 13843124 | 83.7459228 | 13843124 | 83.7459228 |
| B9WC | 12561364 | 10886204 | 86.6641871 | 10886204 | 86.6641871 |
| B9WH | 11374292 | 9312407 | 81.8724102 | 9312407 | 81.8724102 |
| B12CC | 13842452 | 10145419 | 73.2920656 | 10145419 | 73.2920656 |
| B12CH | 15202044 | 10784951 | 70.9440849 | 10784951 | 70.9440849 |
| B12WC | 11239388 | 8883081 | 79.0352731 | 8883081 | 79.0352731 |
| B12WH | 9929494 | 7469932 | 75.2297348 | 7469932 | 75.2297348 |
| B15CC | 19015044 | 16681485 | 87.7278275 | 16681485 | 87.7278275 |
| B15CH | 15472266 | 11259040 | 72.7691729 | 11259040 | 72.7691729 |
| B15WH | 9602548 | 8257182 | 85.9894895 | 8257182 | 85.9894895 |
| B15WC | 23955856 | 20615604 | 86.0566368 | 20615604 | 86.0566368 |
| B18CC | 12739676 | 10029528 | 78.7267117 | 10029528 | 78.7267117 |
| B18CH | 27117408 | 13282380 | 48.9810088 | 13282380 | 48.9810088 |
| B18WH | 16087848 | 13806047 | 85.8166176 | 13806047 | 85.8166176 |
| B18WC | 14948218 | 12180585 | 81.4851978 | 12180585 | 81.4851978 |
| B21CC | 29966274 | 25641706 | 85.5685495 | 25641706 | 85.5685495 |
| B21CH | 25180630 | 20865188 | 82.8620571 | 20865188 | 82.8620571 |
| B21WC | 4849778 | 4323444 | 89.1472558 | 4323444 | 89.1472558 |
| B21WH | 10453152 | 8314876 | 79.5441987 | 8314876 | 79.5441987 |
| C0CC | 67481492 | 56790392 | 84.1569893 | 56790392 | 84.1569893 |
| C0CH | 16174462 | 13366287 | 82.638217 | 13366287 | 82.638217 |
| C0WC | 15650800 | 12283612 | 78.4855215 | 12283612 | 78.4855215 |
| C0WH | 21902796 | 19659714 | 89.7589239 | 19659714 | 89.7589239 |
| C3CC | 11850992 | 9916294 | 83.6748012 | 9916294 | 83.6748012 |
| C3CH | 14785154 | 13211315 | 89.3552749 | 13211315 | 89.3552749 |
| C3WC | 15493128 | 12084275 | 77.9976452 | 12084275 | 77.9976452 |
| C3WH | 347894266 | 274476627 | 78.8965654 | 274476627 | 78.8965654 |
| C6CC | 25785536 | 20910485 | 81.0938543 | 20910485 | 81.0938543 |
| C6CH | 39271560 | 32068817 | 81.659137 | 32068817 | 81.659137 |
| C6WC | 45825694 | 37686733 | 82.2393066 | 37686733 | 82.2393066 |

| | | | | | |
|---|---|---|---|---|---|
| C6WH | 18207706 | 14967677 | 82.2051773 | 14967677 | 82.2051773 |
| C9CC | 38009616 | 31354238 | 82.4902783 | 31354238 | 82.4902783 |
| C9CH | 21017608 | 16743888 | 79.666002 | 16743888 | 79.666002 |
| C9WC | 18405324 | 16022285 | 87.0524474 | 16022285 | 87.0524474 |
| C9WH | 25403406 | 22061130 | 86.8431973 | 22061130 | 86.8431973 |
| C12CC | 7980740 | 6907735 | 86.5550688 | 6907735 | 86.5550688 |
| C12CH | 17272966 | 14140404 | 81.8643654 | 14140404 | 81.8643654 |
| C12WC | 14094022 | 11688528 | 82.9325227 | 11688528 | 82.9325227 |
| C12WH | 9448908 | 7481518 | 79.1786522 | 7481518 | 79.1786522 |
| C15CC | 19734856 | 16039426 | 81.2746037 | 16039426 | 81.2746037 |
| C15CH | 8833696 | 7539416 | 85.3483751 | 7539416 | 85.3483751 |
| C15WC | 6788902 | 5376941 | 79.201924 | 5376941 | 79.201924 |
| C15WH | 45314274 | 34367233 | 75.8419588 | 34367233 | 75.8419588 |
| C18CC | 13891524 | 11369411 | 81.8442311 | 11369411 | 81.8442311 |
| C18CH | 13758634 | 11727538 | 85.2376624 | 11727538 | 85.2376624 |
| C18WC | 22035476 | 17345224 | 78.7149958 | 17345224 | 78.7149958 |
| C18WH | 11407152 | 9863439 | 86.467148 | 9863439 | 86.467148 |
| C21CC | 12990238 | 10520906 | 80.9908641 | 10520906 | 80.9908641 |
| C21CH | 22029706 | 18089278 | 82.1131158 | 18089278 | 82.1131158 |
| C21WC | 18762700 | 15178243 | 80.8958359 | 15178243 | 80.8958359 |
| C21WH | 32922920 | 25711302 | 78.0954484 | 25711302 | 78.0954484 |

**Table S2:** Genes referenced in this manuscript.

| Gene_Name | AGI |
|---|---|
| CCA1 | AT2G46830 |
| LHY | AT1G01060 |
| TOC1 | AT5G61380 |
| PRR5 | AT5G24470 |
| PRR7 | AT5G02810 |
| PRR9 | AT2G46790 |
| LUX | AT3G46640 |
| ELF3 | AT2G25930 |
| ELF4 | AT2G40080 |
| BAM1 | AT3G23920 |
| UBA2A | AT3G56860 |
| UBA2B | AT2G41060 |
| UBA2C | AT3G15010 |
| PUB42 | AT1G68940 |
| PUB55 | AT1G01660 |
| PUB56 | AT1G01670 |
| CSU4 | AT5G63440 |
| BHLH35 | AT5G57150 |
| BGLU17 | AT2G44480 |
| APX2 | AT3G09640 |
| RVE4 | AT5G02840 |
| RVE8 | AT3G09600 |
| LNK1 | AT5G64170 |
| LNK2 | AT3G54500 |
| ERF53 | AT2G20880 |
| ERF54 | AT4G28140 |
| ATPS1 | AT3G22890 |
| ATPS2 | AT1G19920 |
| ATPS3 | AT3G47420 |
| ATPS4 | AT5G43780 |
| APR1 | AT4G04610 |
| APR2 | AT1G62180 |
| APR3 | AT4G21990 |
| SiR | AT5G04590 |

| | |
|---|---|
| OASA1 | AT4G14880 |
| OASA2 | AT3G22460 |
| OASB | AT2G43750 |
| OASC | AT3G59760 |
| HSFA1a | AT4G17750 |
| HSFA1b | AT5G16820 |
| HSFA1d | AT1G32330 |
| HSFA1e | AT3G02990 |
| HSFA2 | AT2G26150 |
| HSFA3 | AT5G03720 |
| HSFA4a | AT4G18880 |
| HSFA4c | AT5G45710 |
| HSFA5 | AT4G13980 |
| HSFA6a | AT5G43840 |
| HSFA6b | AT3G22830 |
| HSFA7a | AT3G51910 |
| HSFa7b | AT3G63350 |
| HSFA8 | AT1G67970 |
| HSFA9 | AT5G54070 |
| HSFB1 | AT4G36990 |
| HSFB2a | AT5G62020 |
| HSFB2b | AT4G11660 |
| HSFB3 | AT2G41690 |
| HSFB4 | AT1G46264 |
| HSFC1 | AT3G24520 |
| TIP2 | AT3G26520 |

125 **Table S3:** Primers used in this manuscript

| Purpose | Primer 1 | Primer 2 |
|---|---|---|
| Genotyping Salk_039895 *bam1* line | TCAACCTCTTCAACAACCCAC | CGCTTAATTTATCGCATCAGC |
| qPCR confirmation of *bam1* mutant upstream of insertion | GGCGAGAATGATGACACCTAAA | GCACGTGTCGCACCTAAA |
| qPCR confirmation of *bam1* mutant downstream of insertion | GGTGACTCTGTCACTATTCCTC | ACTCATGGTTCCTTCTTCCC |
| Amplification of UBA2A promoter | GACTCTGGATCCTTTCGAATTCTAATGG AGATTTTAGGG | ACTGTGAAGCTTAAGTCTCG CGGGAAAC |

126
